# Heterogeneous and higher-order cortical connectivity undergirds efficient, robust and reliable neural codes

**DOI:** 10.1101/2024.03.15.585196

**Authors:** Daniela Egas Santander, Christoph Pokorny, András Ecker, Jānis Lazovskis, Matteo Santoro, Jason P. Smith, Kathryn Hess, Ran Levi, Michael W. Reimann

## Abstract

Simplified models of neural networks have demonstrated the importance of establishing a reasonable tradeoff between memory capacity and fault-tolerance in cortical coding schemes. The intensity of the tradeoff is mediated by the level of neuronal variability. Indeed, increased redundancy in neuronal activity enhances the robustness of the code at the cost of the its efficiency. We hypothesized that the heterogeneous architecture of biological neural networks provides a substrate to regulate this tradeoff, thereby allowing different subpopulations of the same network to optimize for different objectives. To distinguish between subpopulations, we developed a metric based on the mathematical theory of simplicial complexes that captures the complexity of their connectivity, by contrasting its higher-order structure to a random control. To confirm the relevance of our metric we analyzed several openly available connectomes, revealing that they all exhibited wider distributions of simplicial complexity across subpopulations than relevant controls. Using a biologically detailed cortical model and an electron microscopic data set of cortical connectivity with co-registered functional data, we showed that subpopulations with low simplicial complexity exhibit efficient activity. Conversely, subpopulations of high simplicial complexity play a supporting role in boosting the reliability of the network as a whole, softening the robustness-efficiency tradeoff. Crucially, we found that both types of subpopulations can and do coexist within a single connectome in biological neuronal networks, due to the heterogeneity of their connectivity. Our work thus suggests an avenue for resolving seemingly paradoxical previous results that assume homogeneous connectivity.

## 1 Introduction

Neuronal activity is typically unreliable, i.e., highly variable across repetitions ^1–3^. While part of this variability may have a functional role, some of it is simply noise coming from a wide variety of sources, such as synaptic failure ^4–6^. This variability poses a fundamental problem for a robust, i.e., error-resistant, code. One proposed solution is population coding, i.e., encoding the same information across multiple neurons, e.g., through neuronal assemblies^7–9^. If the activity of the neuronal population is highly correlated and its activity manifold^10^ therefore low dimensional, the added redundancy provides a population code that can overcome noise and thus encode information robustly. On the other hand, if the activity is mostly uncorrelated and its activity manifold therefore high dimensional, the information is encoded efficiently^11–16^.

Conceptually, thus, circuits evolve to optimize the tradeoff between *robustness* and *efficiency*, mediated by the degree of neuronal *reliability*. We explain this interaction with the following simple thought experiment. Consider a neural circuit in a sensory region. Assume neurons have a given level of reliability, which measures how similar their responses are across repetitions of a stimulus (Fig. 1A). Note that for the purpose of this illustration we make no claim about the source of variability, e.g., noise or non-stimulus-related information. If the neurons were perfectly reliable, a number of (uncorrelated) properties up to the number of neurons could be encoded without errors, i.e., at maximal robustness (Fig. 1B, pink). For less-than-perfect reliability, this number of properties can be encoded only at the price of a loss of robustness. Alternatively, the level of robustness can be maintained by reducing the number of properties encoded (Fig. 1B, purple). The severity of the tradeoff can be reduced by increasing neuron reliability, thereby shifting to a more favorable curve in the plot.

**Figure 1:**
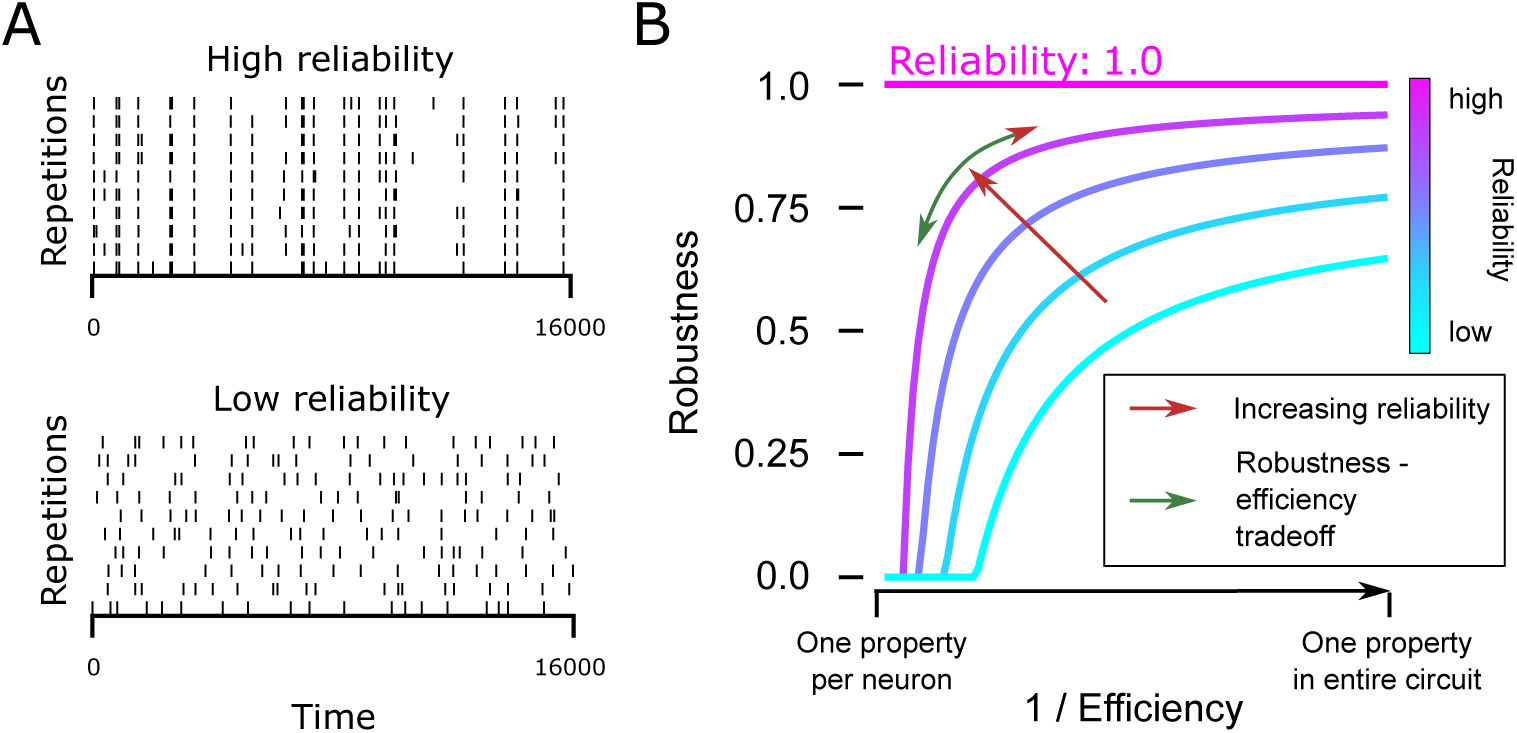
The efficiency - robustness tradeoff. **A:** Examples of neurons with low/high reliability. **B:** There is a tradeoff between efficiency and robustness in a neuronal circuit. The severity of the tradeoff is mediated by the level of neuronal reliability.

Note that related notions of efficient coding have been formulated and mathematically investigated before, two of which appear frequently in the literature: sparse coding^17,18^ and maximal network capacity ^19–21^. Though based on different mathematical foundations, information theory and dynamical systems, respectively, these two notions are not in conflict with our definition. Moreover, they also confront an efficiency-robustness compromise exhibited as a tradeoff between minimizing the number of active neurons while remaining “overcomplete” in the first case, and maximizing both the number of basins of attraction and the distance between them in the second. In both cases, the severity of the tradeoff is mediated by a certain level of amplitude of a noise term playing the role of neuronal reliability. Furthermore, on a related note but from the perspective of complex dynamical systems, the question of what optimizes computational capacity has been explored ^22–25^. In the discussion, we further examine how their concept of “computing at the edge of chaos” or “self organized criticality” connects to our work.

The main purpose of this work is to study how the efficiency-robustness tradeoff plays out in neural circuits. Is it uniform between subnetworks, such as layers? Given that synaptic connectivity shapes the neural code, how does its structure affect the outcome of the tradeoff? Beyond this tradeoff (Fig. 1B, green arrow), increasing reliability globally is a way of improving robustness of the code without loss of efficiency (Fig. 1B, red arrow). Can we find evidence of connectivity mechanisms enhancing reliability of firing? If so, are they employed equally, or do they favor some subpopulations?

As our goal involves linking the structure of connectivity to a functional measure, we need relevant network metrics, as well as structural data together with co-registered functional data. Higher levels of functional redundancy have been achieved by more tightly interconnecting a subpopulation ^26–29^. Thus, we employ a mathematical framework based on finding densely connected motifs called *directed simplices* and describing how they are connected to each other to form larger motifs. A directed simplex of dimension *k* is a motif of *k* + 1 neurons all-to-all connected in feedforward fashion(see Box). Simplices have been demonstrated to have functional relevance^30^. In particular, they are related to the reduction of dimensionality of neural activity and the formation of assemblies, as the correlation of spiking activity of neurons increases with the dimension of simplices they participate in ^31,32^. Structurally, they have been shown to be over-expressed motifs in neural networks at all scales ^33–37^, even though constructing random sparse directed graphs with a specified overexpression of simplices is an unsolved mathematical problem^38,39^. Finally, this framework offers great analytical flexibility as it provides a range of network metrics that describe not only the global structure of a graph, but also its local structure, i.e., node- or even edge-based metrics. These node-based metrics are generally strongly related to other notions of node centrality obtained from pairwise interactions ^33,40^. Thus, they provide a complementary perspective on other metrics describing higher-order structure, such as rich clubs, clustering, or small world coefficients^41–44^.

#### Simplices in directed networks

A directed simplex of dimension *k* is a motif of *k* + 1 neurons where the connectivity between them satisfies the following criterion: there exists a numbering of neurons in the motif from 0 to *k*, such that if *i* < *j* then there exists a synaptic connection from *n_i_* to *n_j_* (Fig. 2A). We call neuron *n*_0_ the *source* and neuron *n_k_* the *sink* of the simplex.

As the functional properties central to this work, i.e., efficiency and robustness, are population-based, we need population-based structural metrics that we can link to them in addition to the node-based ones. Thus, we developed a population-based structural metric that is also related to simplex motifs, which we call *simplicial neighborhood complexity*. Briefly, it compares the synaptic connectivity of a subpopulation to that of a random control.

To demonstrate the usefulness of simplex-based metrics, we first show that they capture non-random trends in many different biological neural networks and that they are related to previously studied phenomena, such as overexpression of reciprocal connections^45,46^. To benchmark our structural metrics, we analyzed several networks with cellular resolution: the mature C. elegans ^47^, drosophila larva ^48^, and an electron-microscopic reconstruction of mouse cortical connectivity ^49^ (MICrONS). Note that, while the MICrONS data set is imperfect, the error rate of this reconstruction has been determined, and we show that the results of our analyses cannot be explained by errors at that rate. Additionally, the MICrONS data set also provides co-registered activity, which we can use to study the structure-function relation. However, the co-registered activity data in MICrONS is too sparse to estimate some population-based functional metrics. Thus, we additionally generated activity data from simulations of a morphologically detailed model^50^ (BBP) to confirm the relevance of the metrics for reliability, robustness and efficiency, and to develop a detailed picture of the structure-function relation.

We believe that the BBP network is relevant for our use case as it provides full certainty and control of both structural and functional results. It enables more complex analyses and connectivity manipulations that allow us to link structure of connectivity to function in more detail and, in some cases, in causal ways. Further, it captures much of the biological detail that is relevant for our field of enquiry, such as sources of variability, in particular, synaptic failure, spontaneous release, and stochastic ion channels. Extrinsic input from other cortical regions is modeled with a relatively simple noise model. Additionally, the BBP model has already been used to study the question of noise and firing variability ^30^. Finally, we will demonstrate that the BBP network reproduces the same non-random simplex-based structural features that are present in biological neural networks. While they emerge less strongly in the model, this is nevertheless highly significant, as we will demonstrate that the features cannot emerge by chance. We will also demonstrate that they correlate with measures of efficiency and reliability in qualitatively the same way as in biological neural networks.

With these tools and data sources, the following are our main results. First, in Section 2.3 we show that the greater the number of simplex motifs to which a neuron belongs, the higher the correlation of its activity with that of the population as a whole. This leads to coding in subnetworks with with higher density of simplex motifs being less efficient, but generally more robust, demonstrating the impact of simplex motifs in positioning the cortical code along the robustness-efficiency tradeoff (Figure 1 green arrow). In particular, the highest level of simplex overexpression leads to low efficiency with little to no increase in robustness. In Section 2.4 we show that the presence of such highly complex subnetworks globally increases spiking reliability of neurons (Figure 1 red arrow). Further, we show that the increase in reliability through higher density of simplex motifs and reciprocal connections also affects reliability at the single neuron level, albeit weakly.

## 2 Results

### 2.1 The overexpression of reciprocity is shaped by the higher-order structure

To confirm previous results about the structural relevance of reciprocal connectivity and simplex motifs (see Box), we counted their overexpression in the following four available connectomes at cellular resolution: First, the electron microscopy reconstructions of the full brain connectivities of the adult *C. elegans* ^47^ and the *Drosophila larva* ^48^. Second, the electron microscopy reconstructions of 1 *mm*^3^ volume of mouse primary visual cortex (*MICrONS* ^49^) and a morphologically detailed model of *∼*2 *mm*^3^ of rat somatosensory cortex (*BBP* ^50^). See Methods for details on the data sources for all connectomes.

Across all connectomes we found an overexpression of reciprocal connections and directed simplex counts when contrasted to relevant control models of the same size (Fig. 2 B, C). The controls considered were: *Erdős–Rényi* controls (ER), *directed configuration model* controls (CM) and *distance dependent* controls (Dist), which preserve: the overall density of connections, the in and out degrees of the nodes, and the density of connections at a given distance, respectively (see Methods). Note in particular, that the CM controls will preserve the known long-tailed degree distributions ^51–54^. The distance dependent controls were only performed for MICrONS and BBP, given that Euclidean distances between cell bodies might not be the most relevant factor determining connection probability for C. elegans and Drosophila. Overexpression of reciprocal connections was weakest in the BBP model connectome, to a degree where it was not overexpressed compared to the distance-dependent control. All other overexpressions of reciprocal connections and simplex counts were statistically significant at the most stringent significance levels. In particular, this shows that distance dependent connectivity and degree heterogeneity contribute but do not fully explain the high complexity of the biological connectivity structure across species. This is unlike in simple network models, optimized for information storage, where higher-order structure can be explained from a wide degree distribution ^46^.

**Figure 2:**
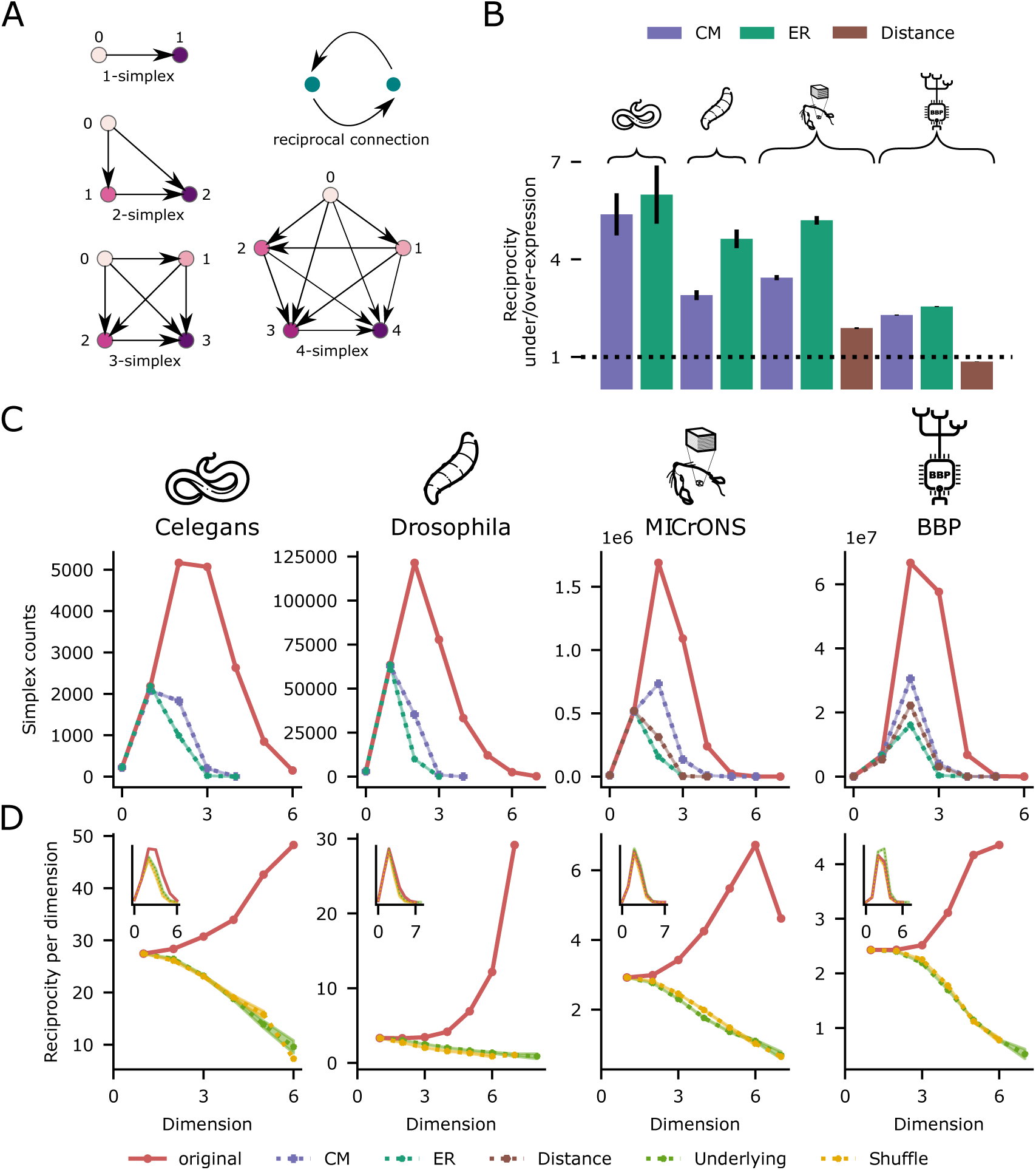
Over expression of directed motifs. **A:** Reciprocal connection (green nodes) and simplices (purple nodes). **B:** Percentage of reciprocal connections in each connectome relative to mean of the percentage of 100 controls of different types: Erdős–Rényi, Configuration model and Distance dependent (see Methods). A value of 1 indicates the original connectome has the same percentage of reciprocal connections than its corresponding control; values above/below 1 represent over/under expression; vertical bars show the standard deviation. All values are significantly different than the mean with p-values under 3 *×* 10*^−^*^1^^55^ for a two sided one sample t-test. **C:** Overexpression of simplex motifs with respect to 10 random controls, which by design match the counts of the original network for dimensions 0 and 1. All counts in dimensions greater than one are significantly higher than the mean of the controls with p-values under 4*×*10*^−^*^18^ for a one sided one sample t-test. **D:** Percentage of reciprocal connections in the subgraph of simplices of each dimension contrasted with the same curve for controls where just the directionality of the connections is modified (see Methods). Inset: Simplex counts of the directional controls are close to the original ones.

In addition to simplex counts as a global metric of network complexity ^55–57^, we can quantify the participation of individual elements in that structure. For example, *k-node participation*, defined as the number of *k*-simplices to which a node belongs, provides a generalized notion of node centrality highlighting the relation between simplex counts and more classical metrics in network science. Indeed, we confirmed that in- and out-hub nodes have higher node participation than the rest of the population (Fig. S1). Moreover, node participation is correlated to closeness centrality, a metric measuring how close a node is to all other nodes in the graph in terms of path distance ^58^. Similar observations, relating simplex counts and other centrality metrics, have been made before for other biological networks ^33,40^.

On the other hand, *k-edge participation*, defined as the number of *k*-simplices an edge is part of, gives a notion of edge centrality that has been observed to be meaningful for describing synaptic strength, and predicting plastic changes, both in MICrONS and the BBP connectome^59^. We can filter edges by the maximum dimension at which this metric is non-zero. Doing this, we found that edges with a high maximum dimension are more likely to be reciprocal and this preference increases with dimension (Fig. 2D, red curves). Moreover, this preference is not merely a combinatorial artifact. To show this, we contrasted these curves with those obtained from two types of random controls: one (*rc-Sh*) only modifies the location of reciprocal connections and the other (*Und*) modifies their location together with the direction of all the connections (see Methods). Crucially, these controls are quite stringent in that they have the same underlying undirected graph as the original and thus approximately preserve the number of simplices across dimensions (Fig. 2D, inset). We observed that the percentage of reciprocal connections in these controls actually decreases with dimension, showing that combinatorially it is less likely for a reciprocal connection to exist on higher dimensional maximal simplices if these where placed at random.

Note that we studied only a central subvolume of the MICrONS connectome, in order to avoid any boundary artifacts, as well as to have a cylindrical volume comparable to the BBP connectome. Furthermore, synaptic contacts for the MICrONS connectome were automatically segmented and their corresponding partners automatically assigned. It has been estimated and reported that the automated detection has a: “precision of 96% and recall of 89% with a partner assignment accuracy of 98%” ^49^. We therefore generated controls to address all these potential sources of error by: removing 4% of the connections, adding (1/0.89) *−* 1 = 12% of connections, and shuffling 2% of the connections, taking into account precision, recall and partner accuracy respectively. We observed that simplex counts and their relation with reciprocal connections remained unchanged (Fig. S3), showing that our metrics are relevant even at the levels of reported inaccuracy of this data source.

In summary, we confirmed previous results that reciprocal connections (a circular motif) and simplices (a feedforward motif) are strongly overexpressed in biological neural networks (BNNs); and that their locations within the networks are strongly related. These trends also hold true for the BBP network. Moreover, these metrics allowed us to show that the complexity of BNNs can not be explained by only distance dependent connectivity, long tailed degree distributions, or connectome reconstruction errors.

### 2.2 Biological neural networks exhibit a large diversity of neighborhoods

In the previous section we showed that BNNs exhibit non-random global structure. It turns out that individual neurons participate in diverse ways in this structure, which could affect their function. We make this clear in two complementary ways. First, on the level of single neurons we exploit the *simplicial structure*, taking averages of functional or structural properties of neurons weighted by the number of maximal simplices in which they participate (see Methods). This analysis reveals the effect of the simplicial centrality of a neuron on its functional properties. Second, on the level subpopulations, we analyze the synaptic *neighborhood structure* of a neuron, given by all neurons directly interacting synaptically with a central one. We define the *neighborhood* of a neuron *v*, as the subnetwork on *v* together with all neurons connected to it (in either direction) and all connections between them and call *v* its *center* (Fig. 3A). One can then apply any network metric to neighborhoods, a particularly simple example being the *neighborhood size*, i.e., the number of constituent nodes, which closely approximates the sum of the in- and out-degrees of its center (see Methods). As expected from previous research^54,53,41,60^, we found across connectomes that the distribution of neighborhood sizes has a longer tail than in corresponding controls, except for CM controls, which fix the node degrees and thus the neighborhood sizes. However, for a plethora of other network metrics, the distributions of their values are also more long tailed than all corresponding controls (see Fig. S4 for examples) including CM, indicating that this difference is not driven by long-tailed node degree distributions alone.

**Figure 3:**
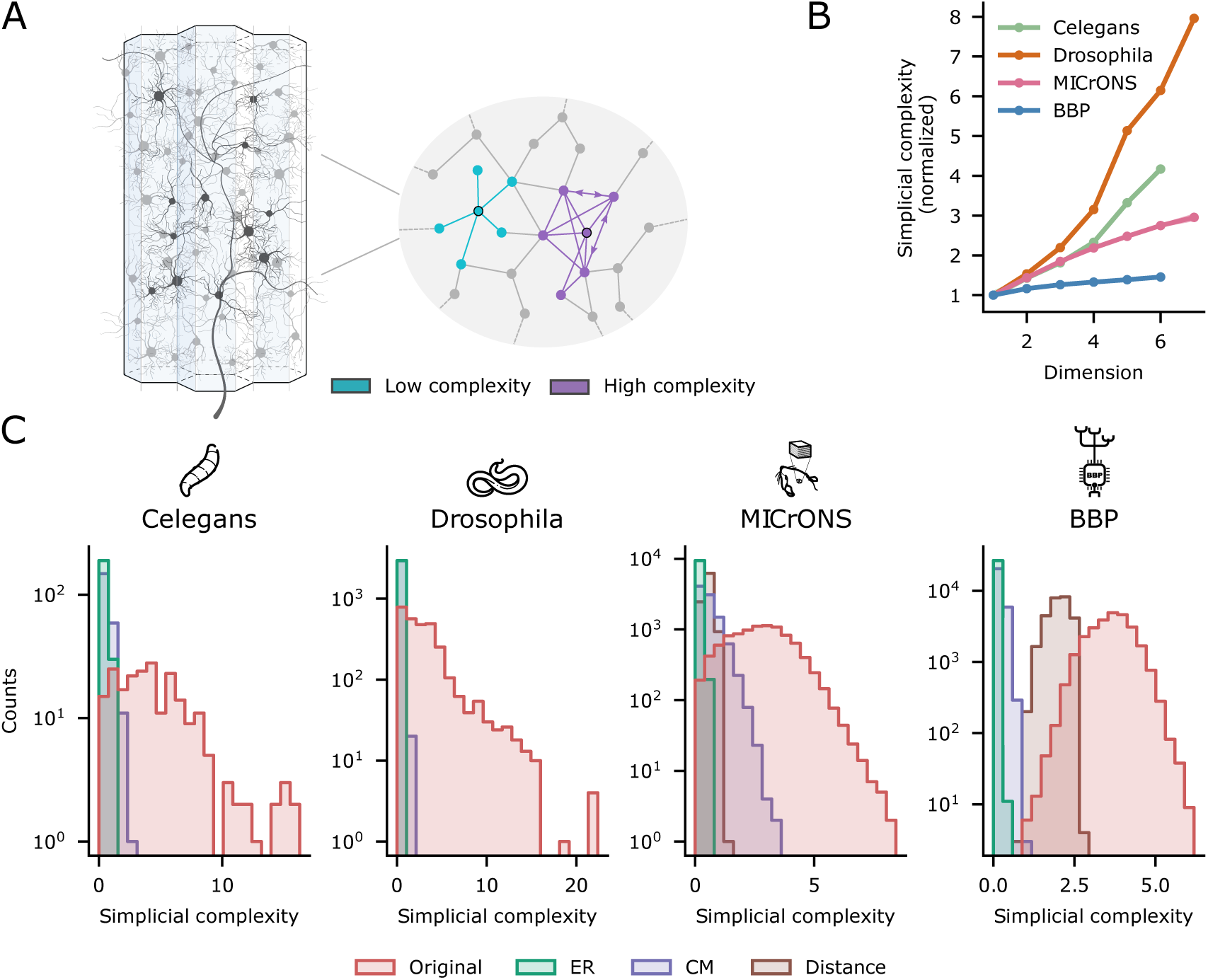
Neighborhood complexity in BNNs. **A:** Illustration showing a connectome and two neighborhoods of high (purple) and low (blue) complexity. **B:** Average of simplicial neighborhood complexity on nodes that are on simplices of a given dimension (see Methods) normalized so that they all have complexity 1 on the first dimension. **C:** In BNNs have longer tailed distributions of neighborhood complexity than relevant controls (described in Figure 2C)

This suggests that the neighborhoods of a BNN exhibit a wide range of complexity, which may fulfill different functional roles. To test this hypothesis, we need to be able to capture this complexity in a single number independent of the neighborhood size. We propose a metric of neighborhood complexity based on its distance to a random network of the same size in terms of normalized simplex counts and call this *simplicial neighborhood complexity* (see Methods and Fig. S5 for an explicit example). We will often refer to this measure as *neighborhood complexity* in short, to ease readability. The distribution of its values is also long-tailed with respect to controls (Fig. 3C). We computed the average neighborhood complexity across maximal simplices and found that it increases with dimension for all connectomes (Fig. 3B); indicating that the simplicial structure and neighborhood complexity are strongly related Additionally, simplicial neighborhood complexity is related to the density of reciprocal connections in the neighborhood (Fig. S6C) as well as to an alternative measure of neighborhood complexity based on distances of degree distributions instead of simplex counts (see Methods; Fig. S6D). The latter confirms the robustness of our metric.

In summary, a structurally highly diverse set of local subnetworks (i.e., neighborhoods) together form the non-random global structure of BNNs. Moreover, the complexity of neighborhoods can be measured by a variety of metrics that are related to each other but complementary. We chose simplex counts as our main metric, as it captures the complexity of the global structure of BNNs and has demonstrated functional relevance.

### 2.3 Low complexity neighborhoods are efficient

So far, we have shown that BNNs exhibit a wide range of neighborhood complexities, as measured by a variety of network metrics. We hypothesize that this heterogeneity provides a substrate for the diversity observed in neuronal function related to the robustness-efficiency tradeoff. More precisely, prior work has shown that the spike correlation between synpatically connected increases with the dimension of the largest simplex of which the connection forms an edge ^31,49^. We therefore hypothesized that on the single cell level, the correlation of the spike train of a neuron with the overall activity of the circuit increases with the number of simplices to which it belongs. On the subpopulation level, neurons belonging to a neighborhood of high simplicial complexity should exhibit more highly correlated activity among themselves. Both effects would reduce the efficiency of the code.

We tested these hypotheses for the MICrONS and BBP connectomes, which have co-registered structural and functional data. For MICrONS, calcium traces of a sparse, subset of neurons (up to 2.5% per scan and up to 20% when pooled across scans; see Methods) were reported for five classes of naturalistic stimuli^49^ (see Methods; Fig. S7A). For BBP, the spike trains of all neurons were reported for eight repeated stimuli (see Methods; Fig. S7B).

To test the first part of the hypothesis, we built on previous work, where a wide spectrum of functional roles was observed experimentally, showing that neurons can differ in their activity correlation to the overall firing of the population, i.e., their *coupling coefficient*, ranging from strongly coupled *choristers* to weakly coupled *soloists* ^61,62^. We found for both MICrONS and BBP that the simplicial structure shapes the value of the coupling coefficient: neurons that belong to higher dimensional maximal simplices have higher coupling coefficients (Fig. 4A). This shows that soloists/choristers are in the lower/higher simplicial dimensions of the connectomes. Moreover, Okun et al. ^61^ also found that the coupling coefficient increases with the amount of synaptic input. Conversely, Brunel ^46^ used simpler network models to predict that neurons with large outdegree are active in smaller assemblies. Intuitively, this would reduce their coupling coefficient and lead to a higher node participation in the source than in the sink position. By showing that sinks of simplices are more coupled than sources, we confirm these findings and extend them by showing that the effect is even stronger if the presynaptic population is highly recurrent.

**Figure 4:**
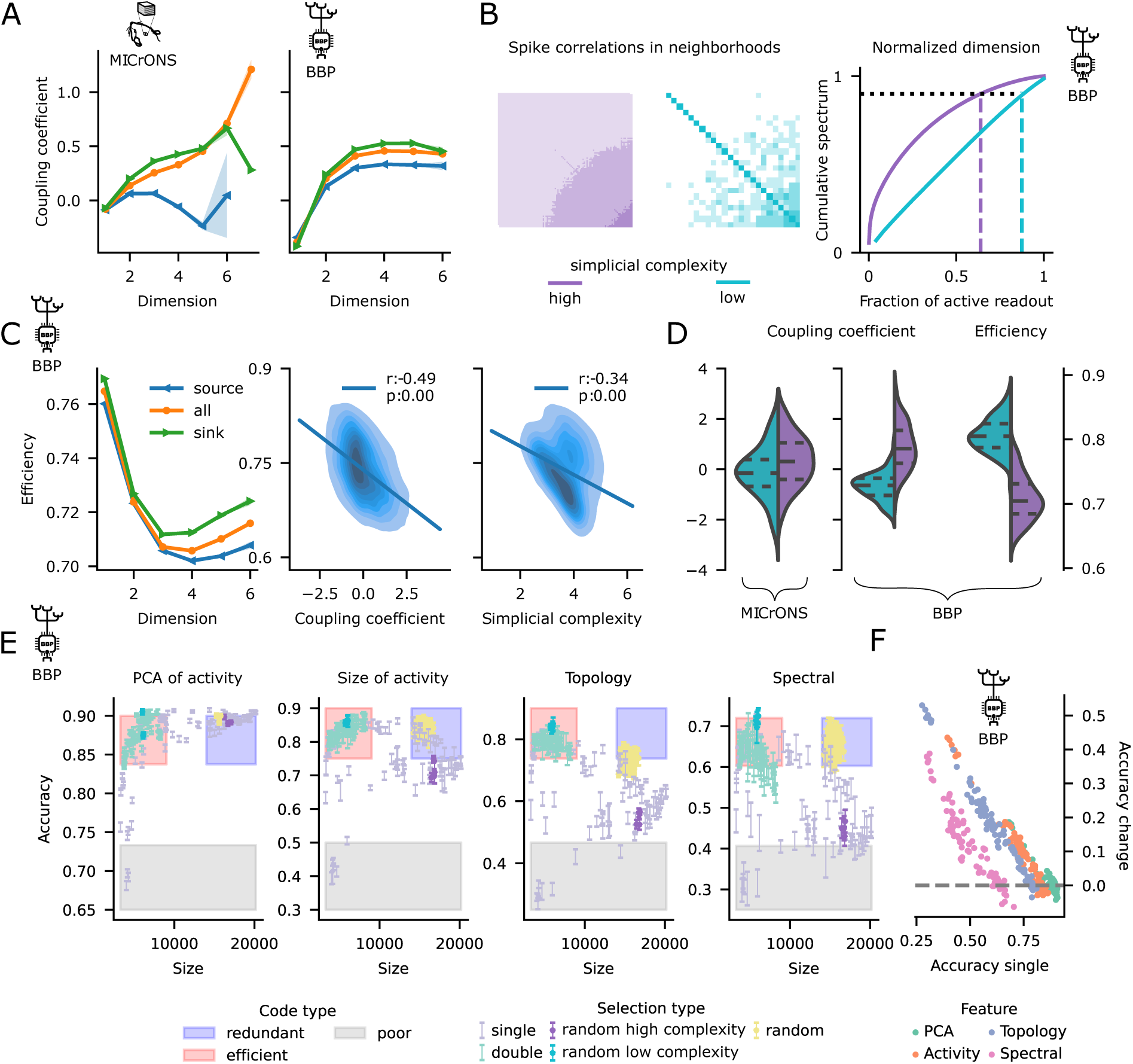
Low complexity neighborhoods are more efficient. **A:** Coupling coefficient (z-scored) increases with maximal simplex dimension for MICrONS (left) and BBP (right). Sinks of neurons (green curves) are more coupled than sources (blue) and the overall population (orange). **B:** Left: correlation matrices of the spike trains of the lowest/highest complexity neighborhoods of the network in the BBP data. Right: their corresponding normalized spectra from which their *efficiency* is determined. **C:** Efficiency for the BBP data, from left to right: Efficiency decreases with maximal simplex dimension colors as in A. Efficiency is anti-correlated to the coupling coefficient, and simplicial neighborhood complexity. **D:** Left: The centers of the 100 least complex neighborhoods have a mean coupling coefficient that is significantly lower than the centers of the 100 most complex ones; with p-values of 1.66 *×* 10*^−^*^3^ and 8.25 *×* 10*^−^*^28^ on the Kruskal–Wallis test for MICrONS and BBP respectively. Right: The opposite effect is observed for Efficiency, with a p-value of 2.60 *×* 10*^−^*^34^ (BBP data only). **E:** Accuracy of the classification task versus size of the subpopulation used for four different featurization methods. Less complex neighborhoods (light blue) consistently give provide higher accuracy with smaller subpopulations; even when these are chosen at random (cyan). **F:** The change in accuracy in the double selection process (see Methods) is anti-correlated to the accuracy of single selection; showing that low complexity is a sufficient condition for efficient coding.

In order to test the second part of our hypothesis, we considered the correlation of activity on the level of subpopulations. Previous work pointed out that mostly uncorrelated activity maximizes the amount of information that can be encoded and is thus a mark of an efficient code ^11–16^. Based on this, we defined the *efficiency* of a neighborhood as the fraction of its neurons required to explain 90% of the variance of its correlation matrix (see Methods, Fig. 4B). The sparsity of the MICrONS functional data set makes it impossible to obtain an estimate of this or any other metric based on pairwise correlations that is representative for neighborhoods of that connectome. Thus, we restrict our analysis to the BBP data set. As we hypothesized, efficiency decreased with the dimension of maximal simplices (Fig. 4C-left) and generally sinks of neurons were more efficient than sources. From a more fine grained perspective, we found that across layers neighborhoods centered on sink neurons were also more efficient than neighborhoods of source neurons and that globally neighborhoods with centers in layer 6 were more efficient than other layers (Fig. S11C). Moreover, the efficiency of a neighborhood is anti-correlated with its simplicial neighborhood complexity, and highly efficient neighborhoods tend to have soloists at their center (Fig. 4C-right). This effect is more pronounced for the *champions*, i.e., the 100 neighborhoods that either minimize or maximize simplicial complexity. Between them we found a statistically significant difference in their efficiency as well as in their coupling coefficient (Fig. 4D-right).

The observed trends should make low complexity neighborhoods efficient readout populations for information processed in a local circuit as well. We tested this in a simple stimulus classification task. For this, we simulated evoked activity in response to a random stream of the 8 stimuli used before (Fig. S7B). Each stimulus was presented 100 times in random order with an inter-stimulus interval of 200 ms. We then tried to determine the stimulus present in a given time window from the spiking activity of a subpopulation of neurons. In order to test the effect of neighborhood complexity, we selected the subpopulations as neighborhoods at different points along the complexity spectrum.

Specifically, we used the neighborhood-based classification pipeline developed in Reimann et al. ^63^ and Conceição et al. ^64^. Briefly, it selects 50 neighborhoods either: at random, or by the ones that maximize or minimize a given network metric from Table S1. We call their original pipeline the *single selection* procedure. This will sample neighborhoods that are either a generic representative of a circuit neighborhood, or structural outliers in one way or another. We explicitly consider outliers, as the values of network metrics over neighborhoods are longtailed ^54,53^ (Fig. S4) and random sampling may miss points from the extreme ends. Then, the spiking activity of the union of neighborhoods is reduced to a set of time series that we call the *feature vectors*. This featurization is done in two ways: either by applying principal component analysis to the binned spike trains (*PCA method* ; bin size 20 ms); or by calculating the value of one of the network metrics listed in Table S1 of the *active subnetwork* of each neighborhood and in each time step (see Methods). That is, we reduce the dimensionality of the spike trains of the subpopulation in different ways to ensure the robustness of our results. Finally, a simple linear classifier is trained to identify the stimulus identity from these feature vectors.

Unsurprisingly, we found that the classification accuracy tended to be higher when the size of the union of the neighborhoods was larger (Fig. 4E, purple bars and Fig. S15).^†^ However, we were primarily interested in efficient coding, not its absolute quality. We considered a neighborhood to be more efficient than another if it reached the same or higher classification accuracy with fewer neurons. As we hypothesized that less complex neighborhoods are more efficient, we executed a *double selection* procedure, where we first selected the 1% least complex neighborhoods and ran the full pipeline described above on those neighborhoods only. When considering the size of the neighborhood union against classification accuracy, we found the following: First, double selection resulted in smaller neighborhoods. Second the data points associated with double selection had comparable or better classification accuracy despite the smaller neighborhoods (Fig. 4E, light blue and cyan bars, red-shaded region). They were therefore more efficient according to our definition.

The overall level of classification accuracy depended on the network metric used to calculate the feature vectors (Fig. S15). For a minority of metrics (9 out of 32) the classification was poor and did not improve for double selection. In these cases, less than *∼* 4.5% (generally less than *∼* 2%) of the entries of the feature vectors where nonzero (Fig. S16). That is, the featurization was degenerate since virtually no input information was given to the classifier. For some selection parameters, classification accuracy was already high in the single selection paradigm. In those cases, it did not further increase for double selection. Instead, double selection ensured high accuracy where accuracy for single selection was low (Fig. 4F). This demonstrates that low complexity is a sufficient condition for efficient coding.

In summary we confirmed our hypothesis that low/high complexity is associated with low/high activity correlations. Consequently, neighborhoods of low complexity encode information most efficiently. On the other hand, neighborhoods of high complexity have choristers at their centers that follow the global activity of the circuit. This is an example of the efficiency-robustness tradeoff.

### 2.4 High complexity neighborhoods promote reliability

As described above, we have shown that the most complex neighborhoods are the least efficient in terms of the amount of information they can encode relative to their size. We have also revealed that the architecture of BNNs contains a diversity of neighborhoods exhibiting long tailed distributions across network metrics. In particular, they exhibit an overexpression of simplex motifs and a larger abundance of complex neighborhoods than simpler architectures (Fig. 2C, S4bottom), leading us to question the purpose of this overexpression. Given that it has been shown that simplex motifs promote network reliability ^30,65^, we hypothesized that one of the functions of the highly complex neighborhoods is to promote *reliability* of the network as a whole, improving the robustness-efficiency tradeoff (Fig. 1B). In order to test our hypothesis of a global effect, we needed to be able to manipulate the overall connectivity and measure the resulting network’s activity. This can naturally only be done *in silico* and thus we started by studying the effect of connectivity on reliability for the BBP connectome. A neuron is reliable if its firing pattern is consistent (i.e., highly correlated to itself) across several equivalent repetitions. The reliability values were computed as the *Gaussian kernel reliability* ^66^ of the spike trains of eight stimuli over ten repetitions (see Methods, and Fig. 1A for two exemplary neurons of high and low reliability).

We employed the recently developed `Connectome-Manipulator` Python framework ^67^, which allows rapid connectome manipulations of network models in the SONATA format ^68^ (see Methods). We performed manipulations that increase or decrease the network’s complexity in different magnitudes while keeping the overall density constant (Fig. S12) and we use overall simplex counts as a metric for global complexity (Fig. 5A).

**Figure 5:**
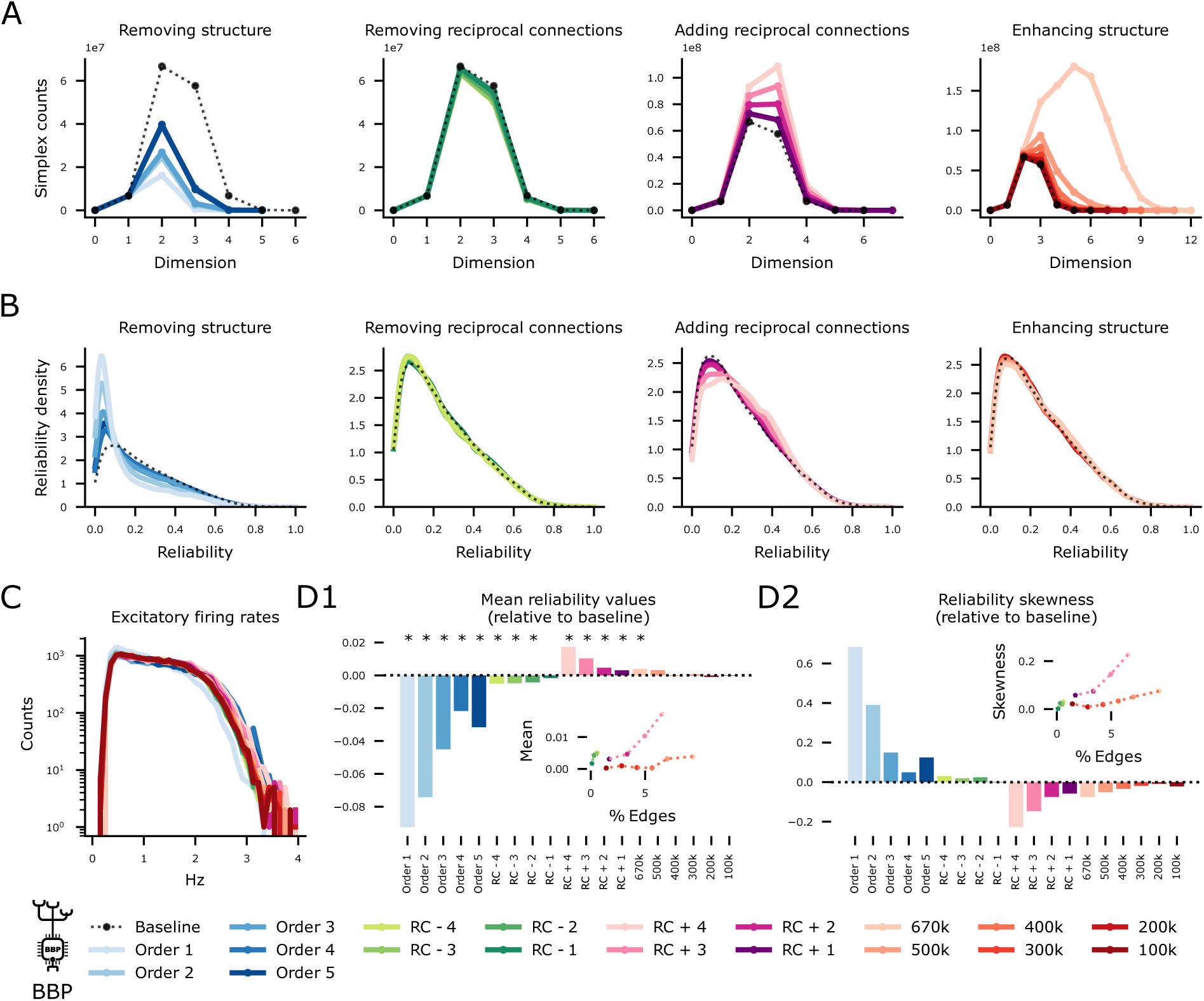
Network complexity enhances network reliability globally. **A:** Simplex counts of manipulated connectomes. Across all panels, black denotes the baseline values, blue and green reduce complexity and red and purple increase complexity; lighter tones denote stronger manipulations than darker tones. **B:** Distribution of the reliability values of all the neurons in the different manipulated connectomes relative to the baseline. **C:** Firing rates remain stable across all manipulations. **D1:** Mean reliability values of the manipulated connectomes relative to the baseline. Positive values denote an increase of reliability in the manipulated connectome, the *∗* marks manipulations for which the change on the mean reliability is statistically significant (with p-values smaller than 0.009 with *n* = 26567 on the Kruskal–Wallis test). Inset, absolute value of the change of the mean versus the % of the edges that have been manipulated. The reduced complexity models are not shown in the inset since virtually 100% of the edges are modified in these manipulations. **D2:** As D1 but for skewness of the distribution rather than the mean value.

We performed four different types of manipulations that *rewire* the model connectome to change its complexity in ways that are consistent with the structural metrics we have shown to shape BNNs across species. That is, they increase or decrease simplex counts while keeping the number of connections fixed or they remove or add reciprocal connections in simplices (see Methods and Table S1). Moreover, different manipulations rewire a different amount of connections. Note however, that achieving a given number of simplex counts and reciprocity in unison remains an open problem in combinatorial and stochastic topology^38,39^.

We found that reducing network complexity always lowered the mean reliability of the network, while the opposite happens for increased complexity (Fig. 5A vs B). At the same time, other spiking statistics, such as firing rate distributions, remained unchanged (Fig. 5C). In most cases the change of mean reliability was statistically significant (Fig. 5D1, KW-test for *n* = 26567), and the p-value as well as the size of the change was strongly determined by the percentage of the edges that had been rewired (Fig. 5D1-inset). On the other hand, even an extreme increase in simplex counts improved mean reliability by less than 1% (Fig. 5D1-light orange), indicating that large simplices are not increasing reliability on their own, but are related to a structural effect that increases reliability and thus provide a convenient way to measure it.

Moreover, the number of connections rewired is not the only driver for the change in reliability; the location of this rewiring within the whole network also plays a significant role. In particular, “adding reciprocal connections” has a stronger effect on the network’s reliability than “enhancing structure” even when a similar number of edges is rewired. Furthermore, this is not due a (slight) increase in overall connection count for the “adding reciprocal connections” manipulation. To demonstrate this, we added or removed the same number of connections as in Figure 5, but instead of doing so in a structured way we do it at random in the whole excitatory network (see Methods and Table S1). There is a substantial difference in the change in reliability between structured and random manipulations (Fig. S13 A, B, C). In the extreme case, where we increased by an 8-fold the number of reciprocal connections, the system has a phase transition from asynchronous to synchronous firing regime (Fig. S13 B2, D1, D2). However, in the random control there is only a moderate increase in the reliability and activity of the network.

Note that even for the most severe reduction of complexity a small number of highly reliable neurons in a long tail of the distribution remained. Therefore, we tested whether the increase in mean reliability was caused by a global shift or rather by increasing the presence of small populations of highly reliable neurons. This can be captured by the skewness of the reliability distributions, which intuitively is given by the normalized difference between the mean and the median (see Methods). A unimodal distribution has a skewness of 0 if it is symmetric and above/below 0 if it is skewed to the right/left. We found that increasing network complexity decreases the skewness of the reliability distribution (Fig. 5D2), showing that the mean reliability increase is due to a global network effect.

Having established that network complexity increases reliability of the network globally, we next studied its potential local effect on neuronal reliability, i.e., on the level of single neurons and neighborhoods. Given that testing local effects does not require connectivity manipulations, we analyzed this for both the MICrONS and BBP connectomes. For MICrONS, reliabilities were provided by the authors of the original publication, for six fixed stimuli that were each presented ten times, as the *Oracle score* of the deconvolved calcium traces over repetitions (see Methods). Note that the definition of the Oracle score is very similar to the Gaussian kernel reliability. We chose to keep the reported values instead of re-computing them, as the authors put a lot of effort into data curation and cleaning that would be difficult to reproduce.

On the single neuron level, we considered reliabilities separately for neurons in individual layers. We found that reliability increased with cortical depth (moderately for the BBP data, slightly for MICrONS) until layer 5, after which it dramatically dropped in layer 6. Furthermore, for both datasets the reliability of a neuron increased with the maximal simplex dimension it participates in (Fig. 6A, orange curves), although this increase was weaker when the baseline reliability in a layer was already high, indicating that reliability cannot be raised over a certain ceiling value. Moreover, in layers 5 and 6, typically associated with cortical-cortical and cortical-thalamic outputs respectively, we found that sources of simplices are more reliable than sinks (see Methods, Fig. 6A blue vs green curves); while the opposite is true for the other layers. These trends were clear in the BBP dataset but were inconclusive in MICrONS. This could be related to the increasing sparsity of the calcium imaging data across layers in MICrONS. Indeed, when pooled across sessions, while up to 35% of the neurons are co-registered in layers 2/3, this drops to 15% and 3% in layers 5 and 6. Within a single scan, the data was too sparse to make a layer by layer split; the general trend of reliability increasing with simplex dimension was still observed, however (Fig. S9).

**Figure 6:**
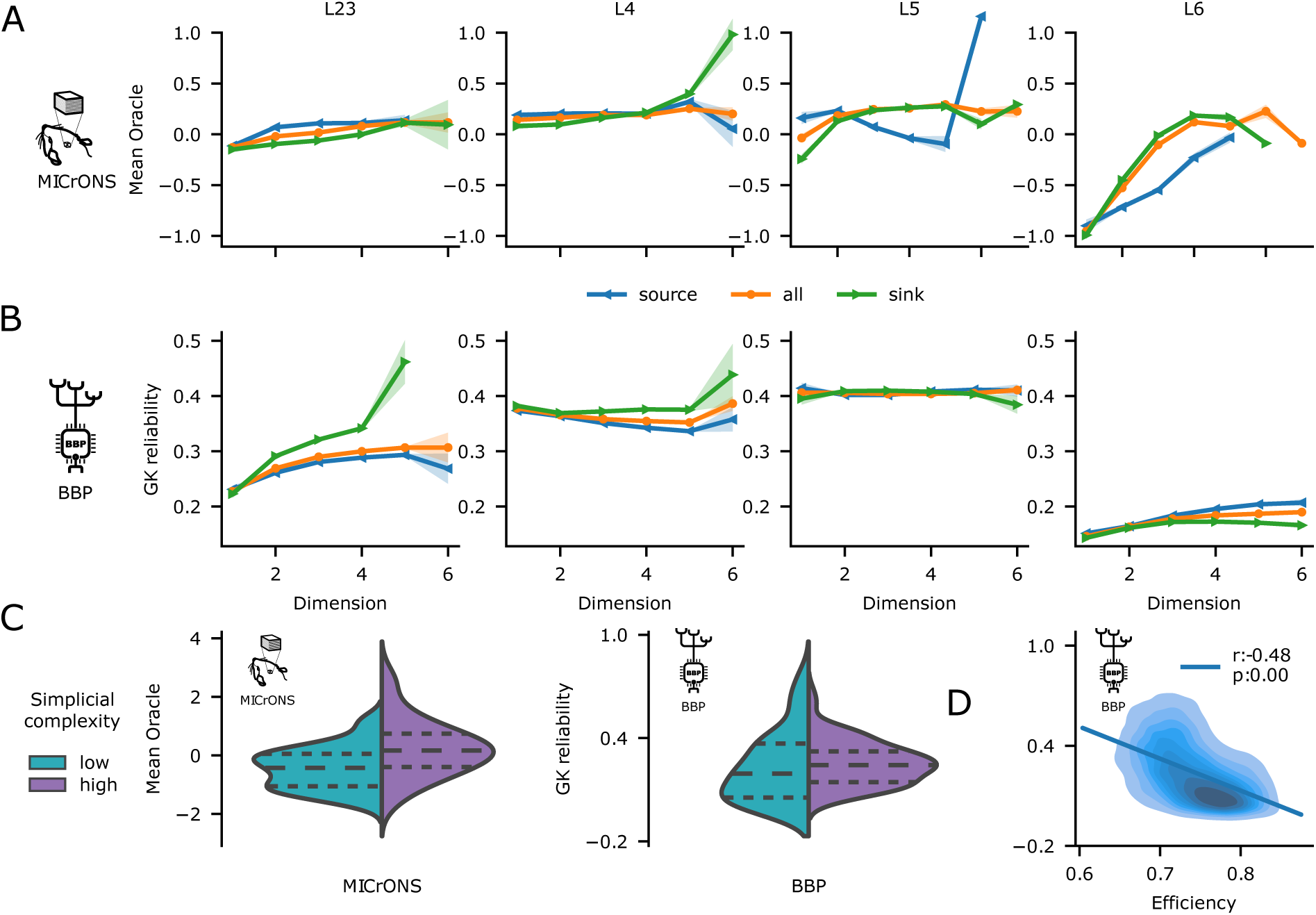
Reliability is shaped by the simplicial structure and network complexity. **A:** Reliability across simplices split by layer for both the MICrONS (Oracle score, top) and BBP (GK reliability, bottom) connectomes. The MICrONS reliability values have been z-scored in order to pool all sessions together maximizing the density of the activity recordings. The roles of the source and target neurons are flipped from the information receiving layers (2-3-4) to the readout layers (5-6). **C:** The centers of the 100 most complex neighborhoods have a mean reliability that is significantly lower than the centers of the 100 least complex ones; with p-values of 3.68 *×* 10*^−^*^7^ and 1.56 *×* 10*^−^*^3^ on the Kruskal–Wallis test for MICrONS and BBP respectively. **D:** The efficiency of a neighborhood is anticorrelated to the neuronal reliability of its center.

On the level of neighborhoods, we contrasted the reliability of individual neurons with their neighborhood complexity for both data sets. Although we found a negligible correlation over the entire connectomes (Fig. S10), we found that between the *champions* i.e., the 100 neighborhoods that either minimize or maximize simplicial complexity, there is a statistically significant difference in their reliability (Fig. 6C). Finally, the efficiency of a neighborhood is anti-correlated to the reliability of its center (Fig. 6D).

In summary, network complexity increases reliability locally in two ways: On the node level, reliability increases with simplex dimension; on the subnetwork level, the most complex neighborhoods have significantly more reliable centers than their least complex counterparts. However, the strongest effect of network complexity is to increase the reliability of the network as a whole. However, this comes at the cost of lowered efficiency in reliability-promoting neighborhoods. The picture that emerges is one of low complexity neighborhoods encoding information highly efficiently, but also robustly due to their interconnections with high complexity neighborhoods that increase the reliability of the network as a whole.

## 3 Discussion

We have investigated the role of the immense diversity of structure found in neuronal networks of different species at cellular resolution. While the presence of wide, long-tailed degree distributions was already known, we found similarly long-tailed distributions of a variety of network metrics, in particular those capturing participation in the higher-order structure of the network. This persisted even in the face of normalization with respect to strong control models. Further, we found that participation of neurons in the higher-order structure was correlated with previously characterized overexpression of bidirectional connectivity^45,61^. This gives rise to subnetworks centered on individual neurons (i.e., neighborhoods) that differ greatly along a spectrum of topological complexity. We linked this spectrum of complexity to circuit function, demonstrating that less complex subnetworks encode information with little redundancy and hence, efficiently. Going beyond abstract theory, we confirmed the efficiency of the subpopulations in a simulated stimulus classification experiment.

To investigate the role of more complex subnetworks, we ran simulations where we increased or decreased the degree of network complexity. The need to study the structure-function relation with connectome manipulations has been pointed out before in Reid et al. ^69^. We found that manipulations changed the mean reliability significantly on a global (whole-network) level, increasing as complexity increased. Additionally, for a given rewiring algorithm, the amount of change is dependent on the fraction of connections rewired. The strength of the effect was stronger for algorithms that decreased complexity, demonstrating that it is easier to destroy than to build the intricate wiring characterizing cortical circuits. In summary, more complex subnetworks promote the robustness of the code and play a supporting role by globally increasing the reliability of the neuron responses in evoked activity.

The contrast between reliability-enhancing and efficiently-coding neurons is related to the previously characterized contrast between “soloists” and “choristers” ^62,61^. We indeed confirmed their results relating connectivity and population coupling, and extended them to demonstrate the importance of higher-order structures. The importance of local connectivity for promoting reliable responses has also been observed before ^30^, but we were able to characterize connectivity motifs that promote reliability. Similarly, functional cell assemblies, which provide robust coding through redundancy, have been demonstrated to emerge through fewer synaptic connections in networks with complex higher-order structure ^32^. This is crucial for BNNs, because unlike artificial neural networks, they are tightly constrained in terms of their connection count: the immensely dense packing of neuronal wiring shows that a given function must be implemented with the least number of synapses possible ^70^. In summary, we confirm, extend and link several known structural and functional features of biological neuronal networks into a single, coherent and quantitative theory. The crucial aspect linking these features is consideration of the higher-order structure of the network, i.e., going beyond pairwise interactions.

Note that there are other notions of efficient coding, two of which are extensive in the literature. First, coming from information theory, the “efficient coding hypothesis”, proposed originally by Attneave ^17^ and Barlow ^18^, links the statistics of natural stimuli to those of the neuronal response. Since natural stimuli are sparse, this theory proposes that an efficient code must also be sparse in the sense that it minimizes the number of active neurons, while preserving the amount of information (in an information theoretical sense). Second, a notion coming from dynamical systems, which posits that information is stored in a network in the form of disjoint basins of attraction in the space of network states, called patterns^19–21^. The “network’s capacity” is the maximal number of patterns it can store and a network is efficient when it has or it is close to maximal capacity. While these two notions of efficient coding have vastly different theoretical foundations, they also exhibit an efficiency-robustness tradeoff: The first, between minimizing the number of active neurons while still remaining “overcomplete” (i.e., containing enough features to encode any stimulus arbitrarily well). The second, between the number of patterns that can be stored and the size of the basins of attraction, which determines robustness against noise.

Interestingly, theoreticians have found a class of networks that are close to maximal capacity, where reciprocal connections and higher-order motifs are overexpressed^46^. However, in contrast to BNNs, they can explain this overexpression with the wide distributions of out-degrees; we speculate that BNNs are a specific subclass of such networks. Furthermore, Iatropoulos et al. ^71^ explored a plasticity rule that explicitly improves robustness in simplified networks and contrasts it with one that maximizes network capacity. They found that the first rule leads to higher density of connections, while the latter leads to lower density. This is consistent with our results, given that higher density networks also have higher simplex counts. We extend the finding beyond connection density to simplex counts, which can be increased to some degree independently of connection density. Additionally, we show that both versions, i.e., high and dense subnetworks, can and do coexist in BNNs, as they are characterized by complex and heterogeneous connectivity. In summary, we also confirm, extend and relate several predictions and theories for simplified neural networks.

On the other hand, what makes a network computationally powerful has also been explored through the lens of complex dynamical systems. It has been observed that maximal computational capacity arises at “the edge of chaos” ^22,23^. That is, at the boundary where the system transitions from ordered to chaotic dynamics. Moreover, it has been observed that the brain might be able to tune itself and evolve towards this region in what is known as self-organized criticality (SOC) ^25^; though it remains unclear whether operating at the edge of chaos drives high performance or is a byproduct of the brain’s need to remain functional^72^. Nonetheless, it has been shown that the brain’s architecture influences the shape of the criticality boundary. Legenstein and Maass ^23^ found that in biologically detailed microcircuits with distance dependent connectivity, the critical regions are determined by the connectivity strength and the level of heterogeneity of connection probability. However, Tadić and Melnik ^24^ and Kilic and Taylor ^73^ found that not only the pairwise connectivity structure but also the higher order structure of the network or its activity are a driving force for the system achieving SOC. In Markram et al. ^50^ it was shown that the BBP network we used in this work is at the boundary of criticality. It was furthermore confirmed for this network that the presence of higher-order structure affects the transition from asynchronous to synchronous activity^30^ and that neuronal reliability increases towards the transition boundary ^74^. This provides a link between the concepts explored in this work and the work on SOC. Interestingly, when we increased by an 8-fold the number of reciprocal connections along simplices the system had a phase transition from asynchronous to synchronous activity, but it did not do so in the random control (Fig. S13 B2, D1, D2). This further shows that indeed critical regions depend on the higher order structure and not only on the pairwise interactions. Our findings suggest new possibilities for future research into the connection between attaining SOC and neighborhood simplicial complexity.

Beyond the general ideas outlined above, we also revealed more specific results relating to the roles of cortical layers that align with their known roles as inputs or output. Our notion of higher-order interactions is based on the presence of simplices, which are motifs with strong directionality, i.e., they have an input or *source* side and an output or *sink* side. In the simulation data we found that, for neurons in layers 2, 3 and 4, efficiency decreased and reliability increased from the source to the sink of a simplex, while in layer 6 this was reversed. Neurons in layer 5 behaved as layer 6 with respect to efficiency, and the picture was non-conclusive for reliability, although they were overall the most reliable population.

The *in vivo* data was more sparsely sampled than the simulated data, leading to noisier results. In fact, the data were too sparse to estimate efficiency or to analyze reliability separately for individual layers within a given recording session. However, the per-layer analysis of reliability could be performed when pooling data across sessions, though the results where not conclusive, as the number of sample neurons decreased towards the deeper layers. Layer 5 is associated with extensive intratelencephalic, i.e., cortico-cortical outputs and layer 6 with cortico-thalamic outputs^75,76^. Therefore, our interpretation is as follows: After inputs into a circuit arrive canonically in layer 4, local connectivity structure is optimized to enhance reliability while progressing activity forward. Within output layers, simplicial structure still increases reliability and enhances efficiency specifically of the output neuron population.

This view is compatible with the theory of “reservoir computing” or “liquid state machines” ^77^, which, in the context of neural circuits, posits that circuit inputs are first projected into a high-dimensional activity state and from there projected to a lower-dimensional readout. Seen in this context, we characterized connectivity mechanisms that increase the robustness of the higher-dimensional projection (ironically reducing the dimensionality of that state), and increase the amount of information contained in the readout. In summary, our hypothesis predicts increasing robustness as the goal for the internal representation of a local circuit, and increasing efficiency as the goal for its output, with specific structures of connectivity associated with both goals and embedded into a single network that promotes the reliability of the whole system. Our work can therefore be used in the future to link also this theory to the topics of non-random connectivity, efficiency and soloist/chorister spiking dynamics.

More specific neuronal roles could be studied when considering larger volumes or full brain connectomes with co-registered activity. Although this data is not yet available at the microscale, there is promising potential for it to become accessible for small animals in the future. On the other hand, human connectomes at the meso-scale have been studied with topological methods before ^33,34,36^ and have been found to have even more complex architectures, e.g., human brain connectomes have been found to have undirected simplices of up to 16 and 20 dimensions, in networks of only 1115 nodes representing different brain regions^34^. Furthermore, their higher order structure has been found to have functional relevance ^33,78–80^. Interestingly, Tadić et al. ^34^ does not only count undirected simplices but also how they are embedded in the full network via what is known as *q*-analysis. In short, in each dimension, one can divide all undirected simplices of dimension *q* into classes, where all the simplices in one class are those where one may travel from one simplex to another by passing through lower dimensional ones. The number of classes in each dimension are called the *number of q-connected components* and are a way of succinctly describing the higher order structure of the network. The theory of *q*-analysis was extended to directed networks in Riihimäki ^81^. This opens a compelling avenue for future research, using *q*-analysis of the level of neighborhoods and its impact on the efficiency-robustness tradeoff. Notably, neighborhoods with high simplicial complexity maximize the number and dimension of simplices in a highly nested and difficult-to-quantify manner. We conjecture however, that once normalized by size, these neighborhoods minimize the number of *q*-components in each dimension, providing a means to quantify their intricate nested structures.

Our view sees trial to trial variability as a problem that needs to be overcome for reliable cortical function. This appears to be in contrast to existing theories where variability explicitly encodes uncertainty about a stimulus^82,83^. However, we believe the views are compatible. First, the theoretical work sees uncertainty encoded primarily on the population level^83^, while we measured reliability on the single cell level. Second, we only investigated the prevalence of variability, not its structure and how it relates to stimuli. Finally, we found remaining uncertainty in the responses of even the most reliable cells. Furthermore, our results demonstrate a wide spectrum of reliability values, and that the mechanisms to reduce it are not uniformly deployed. It is therefore possible that they are deployed in ways that implement the theories of explicit encoding of uncertainty. Moreover, we do not study the effect of correlations between neurons of their trial to trial variability, i.e., of noise correlation, on the network’s efficiency. A common theme on this body of work is that noise correlations are not independent of the network structure ^84^. Investigating how the architecture of BNNs shapes uncertainty encoding and the correlational structure of noise would constitute interesting follow-up work.

Many of our conclusions were drawn from the analysis of electron-microscopic connectomes, primarily the MICrONS data ^49^, which while reconstructed with boundary-pushing methods is still imperfect. However, we have demonstrated that our network metrics are robust against reconstruction mistakes at the reported error rates. Our conclusions relating to efficiency of coding are based on simulations of a morphologically detailed model of cortical connectivity ^50^. They could be confirmed in the future when more or more densely sampled electron-microscopic connectomes with co-registered activity data become available. In the meantime, the qualitative agreement between MICrONS and BBP data in most characterized trends indicates the validity of our model predictions. Quantitatively, non-random trends were slightly weaker in the BBP model than in the MICrONS data, which indicates that the effects observed in the *in silico* model might be even stronger in biology.

While in biology synaptic connections are associated with weights that are lognormally distributed, we considered exclusively unweighted network metrics. A standard procedure in topological data analysis to turn an unweighted metric into a weighted one is to compute the unweighted version repeatedly while filtering the network with a sliding weight threshold. There is a promising future avenue of research, extending our metrics to weighted metrics, enabling investigators to disentangle the the effect of the low versus high strength of synaptic connections. However, such an analysis would require measuring the synaptic strength of millions of connections, which is at the moment not biologically feasible.

Experimental research has demonstrated several ways in which neuronal connectivity is nonrandom with significant higher-order structure ^41,51,33,35,37^. Moreover, small models with simple dynamics have shown the relevance of higher-order motifs in the observed activity ^85–87^ while *in silico* research has shown that the complexity is largely driven by neuronal morphologies ^88–90^. At the same time, in the field of computational neuroscience, the bulk of research is conducted in models with simple connectivity based mostly on pairwise statistics that do not capture the entire complexity of the structure. This is in part due to the difficulty of building such complex networks as well as the need for a framework to quantitatively and systematically describe the higher-order structure. Both are required to demonstrate how this structure matters for the function of a neural circuit, if at all. Our work provides a theoretical framework, based on counting simplex motifs, that links and unifies the diverse aspects of non-random higher-order structure that have been found before. In contrast to global metrics of network complexity (e.g., small-world coefficient or assortativity) simplex counts also provide local metrics of node and edge centrality. These can be used to link functional properties of the neurons to their location in the network (e.g., Fig. 4A, B). Moreover, possibly due to their feedforward nature, simplices have been shown to be functionally meaningful ^31,32,91,59^.

Other motifs have been studied or counted before. Song et al. ^92^, and Brunel ^46^ count all motifs on 3 and up to 5 nodes respectively. Parmelee et al. ^86^ and Curto et al. ^87^, define other types of functionally meaningful motifs which they call “core motifs” and “robust motifs” respectively. In contrast to these, counting simplices can be done more systematically and has been implemented efficiently^93^. On the other hand counting all motifs on *n* nodes present in a network of *N* nodes can only be done for small *n* due to a combinatorial explosion. Indeed, there are 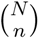 subnetworks on *n* nodes that need to be classified into 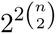 possible network types and further classified into motifs. Similarly, even listing core or robust motifs concretely above dimension 5/6 remains an open problem. Finally, *n*-cycles (i.e., cyclic motifs on *n*-nodes) have similar functional and generalization properties across dimensions as *n*-simplices. However, whilst counting both have the same worst-case complexity ^94^, in practice simplex counts can be computed more efficiently, particularly in large, sparse networks.

Overall, our results and the cited previous research demonstrate that the complex architecture of BNNs provides a substrate to regulate the efficiency-robustness tradeoff, through the existence interconnected low- and high-density subpopulations within its heterogeneous connectivity.

## 4 Limitations of the study

While simplex counts have been efficiently implemented, they can still be computationally prohibitive for certain networks. Heuristically, the computation is harder for larger, denser networks or those that have long fat tail degree distributions. It is also slowed significantly by high degree nodes with high degree neighbors, thus computation time is highly related to the existence of a rich-club in the network. However, the complexity of the computation can not be clearly estimated a priori, since one can not predict the higher order structure just from pairwise metrics. For example, while we were able to count simplices in a network of 4.2 million nodes and 2 billion edges modelling the full somatosensory cortex of a rat’s brain ^90^, counting the number of simplices on an EM reconstruction of a central region of the adult Drosophila brain, a network of 25, 288 nodes and 3.7 million edges ^95^, remains prohibitive due to the highly nested structure of its subnetworks. In the advent of larger and larger data sets becoming available, future work should address how to modify these methods or simplify the connectivity networks in order to address this limitation. These modification could be motivated from the functional-structural implications found in this work in the micro-scale.

## 5 STAR★Methods

### 5.1 Key resource table

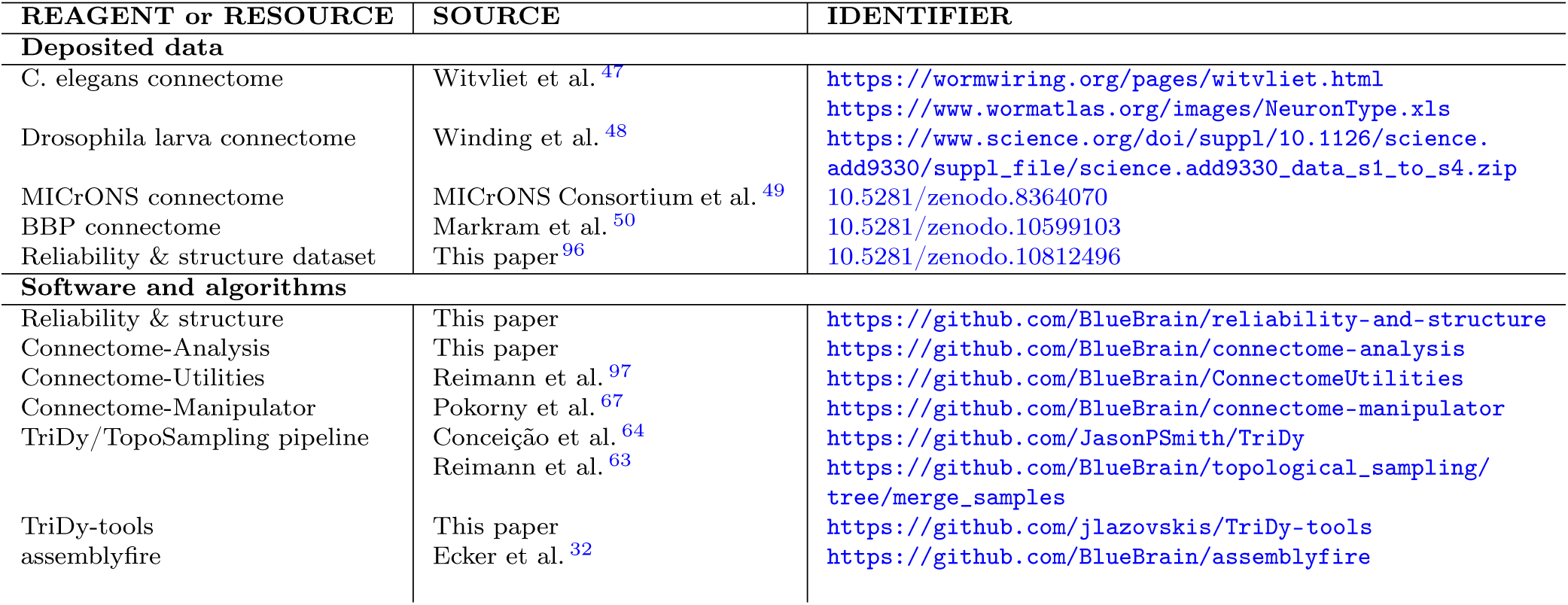

### 5.2 Method details

#### 5.2.1 Structural data sets

We analyzed the connectivity in the following four openly available data sets recorded at cellular resolution. We used the `ConnectomeUtilities` Python package ^97^ to turn these data sets into the common “ConnectivityMatrix” representation format for further analysis.

##### C. elegans ^47^

Electron microscopy reconstruction of the full brain connectivity of the adult C. elegans (Dataset 8; last stage). The data set consists of 324 neurons and 2186 connections formed by 7970 chemical synapses. We did not include electrical synapses in our analyses. Additionally, we discarded 105 isolated neurons of the connectome, i.e., neurons without incoming or outgoing connections.

##### Drosophila larva^48^

Electron microscopy reconstruction of the full brain connectivity of the Drosophila larva. We included both hemispheres but only considered “axo-dendritic” connections. The resulting data set consists of 2956 neurons and 63,545 connections formed by 234,958 synapses.

##### MICrONS^49^

Electron microscopy reconstruction of a cubic millimeter volume spanning all 6 layers of mouse primary visual cortex and three higher visual areas. A reformatted version for easy use with `ConnectomeUtilities` is available on Zenodo ^98^. We excluded the boundaries and restricted the data set to excitatory types. The resulting connectome consists of 9619 neurons and 520,477 connections formed by 610,261 synapses.

##### BBP^50^

Morphologically detailed model of the rat somatosensory cortex ^50^, consisting of more than 210k biophysically detailed neurons belonging to 55 different morphological and 207 different morpho-electrical types. The neurons are connected by over 400M synapses with probabilistic transmitter release and six distinct forms of short-term dynamics of depression and facilitation. The whole circuit can be divided into seven hexagonal columns. To avoid any edge effects, we restricted all analyses (connectivity and activity) to neurons within the central hexagonal column. This data set is available on Zenodo^99^ and can be loaded with `ConnectomeUtilities`. Within that column, we only considered the sub-circuit of 6,717,001 connections between 26,567 excitatory neurons. In the context of connectome manipulations (see Section 5.2.8), we refer to this data set as *baseline connectome*.

#### 5.2.2 Activity data set for MICrONS

We analyzed functional data from the MICrONS dataset^49^. The open source dataset contains calcium traces and deconvolved spike trains of a sparse set of neurons, which are co-registered to the structural dataset. We analyzed data from eight recording sessions from the v661 version of the “minnie65_public” release. The sessions were selected based on two criteria: at least 1000 neurons were scanned and at least 85% of them were co-registered to the structural data. Neurons with a non-unique identifier (2.8% neurons/session on average) were filtered out. Averaged across sessions, 2% of the neurons in the connectome (243 neurons/session) had activity data coregistered to them. Moreover, as calcium imaging is biased towards the superficial layers, the layer-wise distribution of co-registered cells is not uniform, but as follows: layer 23: 36.5%, layer 4: 44.9%, layer 5: 15.3%, and layer 6: 3.3%. Spike trains of neurons restricted to the structural connectome outlined above (243 neurons/session on average, 1817 unique neurons in total, 2% and 15% of all cells respectively) were used to calculate coupling coefficients (see below). As the dataset already contained oracle scores (see below) we did not calculate reliability from the raw spike trains.

#### 5.2.3 Activity data set for BBP

We ran circuit simulations of the morphologically detailed model of the rat somatosensory cortex ^50^, which was converted to the SONATA format ^68^ in order to create manipulated versions of its connectome (see Section 5.2.8). We simulated the entire circuit (original and manipulated versions) using the CoreNEURON simulator ^100^ and recorded the spiking activity of the network in response to a series of stimuli applied to the thalamocortical input fibers. As in Reimann et al. ^63^, the 2170 input fibers were partitioned into 100 clusters of adjacent fibers. Ten such clusters were randomly drawn to form a spatial stimulus pattern. For each cluster belonging to such a pattern, a spike train with a firing rate of 75 Hz was independently generated by an adapting Markov process (with 100 ms adaptation time constant) for 75 ms, followed by a 125 ms blank period with 0.5 Hz firing. We used a random sequence of eight such spatial patterns in two simulation protocols, results of which are available on Zenodo^96^:

##### Reliability protocol

We applied a total number of 80 stimuli (i.e., 10 repetitions per pattern) during 16 s. We reran the experiment 10 times with different simulator seeds using the exact same (random) sequence of stimulus patterns and input spike trains.

##### Classification protocol

We applied a total number of 800 stimuli (i.e., 100 repetitions per pattern) during 160 s. We used a single simulator seed without repeating the experiment.

#### 5.2.4 Random networks

This section contains definitions of the different random connectivity control models and how these were generated. A summary of these is listed in Table 1.

**Table 1:** Types of random graphs. The lower a model is in the table the closer it is to the original graph.

| Label | Model name | Property maintained |
| --- | --- | --- |
| ER | Erdős-Rényi | Overall density of connections |
| CM | Directed configuration model | Sequence of in-/out- degrees |
| Dist | Distance dependent | Density of connection at a given distance |
| Und | Underlying | Underlying undirected graph |
| rcSh | rc-Shuffled | Subgraph of non-reciprocal connections |

The first three controls of Table 1 modify the global structure of the network whilst keeping one property fixed. They are listed in order of complexity. The *Erdős–Rényi* controls (ER) are random directed networks, where each edge is added independently at random with a fixed probability *p* given by 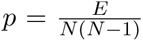, where *E* and *N* are the number of edges and nodes in the original network, thus keeping the global connection probability fixed.

The *directed configuration model* controls (CM) are random networks with (approximately) fixed in- and out-degree sequences, i.e., the vectors whose entries are the in- and out-degrees of all the nodes in the network. To construct such controls we wrote the original network in coordinate format i.e., we wrote its edges as two vectors *sources* and *targets*, such that each pair (*sources*[*i*]*, targets*[*i*]) was a directed edge in the original network. We then shuffled the entries of both vectors independently, which gave a new directed network, with the same degree sequence as the original. Note that this new directed network might contain loops (edges of the type (*a, a*)) or parallel edges (multiple edges with the same source and target). We call these degenerate edges and we removed them. The effect of this removal is that this construction only approximated the original degree sequence. However, the density of degenerate edges tends to decrease as the number of nodes increases ^101^, so the approximation was close. Indeed, while on average *∼* 5% of the edges were lost for the CM controls of C. elegans, only 1.08%, 1.15% and 0.875% of the edges were lost for Drosophila, MICrONS and the BBP data sets respectively.

The *distance dependent model* controls (Dist) are random networks where the edges are added independently at random with a probability that is exponentially decreasing with distance. More precisely, for any pair of neurons, let *d* be the Euclidean distance between the position of their cell bodies. Then, their probability of connection is given by

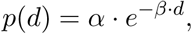

where *α* (the probability at distance zero) and *β* (the decay constant) were determined from the connectivity of the original graph. In order to do so, we computed the pairwise distance between all pairs of nodes in the network and binned them into groups according to this distance. The connection probabilities were estimated in each bin by dividing the number of existing edges by the number of total pairs in that bin. The model coefficients were then determined by fitting the exponential probability function *p*(*d*) to these data points. We did this only for the MICrONS and BBP data sets, obtaining *α ≈* 0.067, *β ≈* 1.18 *×* 10*^−^*^2^*µm^−^*^1^ and *α ≈* 0.111*, β ≈* 7.05 *×* 10*^−^*^3^*µm^−^*^1^ respectively. For C. elegans and Drosophila the Euclidean distances between cell bodies might not be the most relevant factor determining connection probability.

The last two controls in Table 1 are used to control for the possible location of reciprocal connections, while keeping the number of simplex counts close to those of the original graph (specially for large sparse graphs). The *Underlying* controls (Und) are random networks with the same underlying undirected graph as the original and the same number of reciprocal connections. To generate such controls, we first extracted the underlying undirected graph of the original network i.e., the undirected graph with the same nodes as the original and an undirected edge between any two pair of nodes that are connected by a reciprocal or unidirectional edge. We then chose an ordering of the nodes and oriented all edges of the undirected network to go from a smaller to a larger node. Finally, we added the same number of reciprocal connections as in the original graph at random.

The last control, the *rc-Shuffled* control (rcSh) is the tightest control of all. These are random networks where all of the structure of the original network is kept, except the location of the reciprocal connections. To build such a control, for every reciprocal edge in the original network we chose one direction to delete at random. Then we introduced back at random the same number of reciprocal connections along the existing edges.

#### 5.2.5 Network metrics

In a network *G* the *in-degree* and *out-degree* of a node *v* is the number of incoming and outgoing connections from *v*, respectively, and the *total degree* of *v* is the sum of both. We call the *reciprocal degree* of *v* the number of reciprocal connections on *v*, i.e., the number of nodes in *G* that that map to and from *v*.

An *n-simplex* in *G* is a set of *n*+1 nodes which are all to all connected in feedforward fashion. That is, there is a numbering of the nodes 0, 1*,… n* such that whenever *i < j* there is an edge from *i* to *j* in the network *G*. We call *n* the *dimension* of the simplex, the numbering 0, 1*,… n* the *position* of the nodes in the simplex and the edges

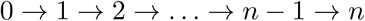

the *spine* of the the simplex. A simplex is *maximal* if its nodes and edges are not contained in a higher dimensional simplex.

We weight the edges of *G* with the maximal dimension amongst all the simplices it participates in and we denote by *G_k_* the subgraph of *G* on the edges with weight *≥ k*. That is, the edges in *G_k_* belong to at least one simplex of dimension *k*. In particular *G*_1_ = *G*, since every edge is a 1-simplex. We then compute the *reciprocity of G at dimension k*, denoted rc*_k_*, as the percentage of reciprocal edges of *G_k_*

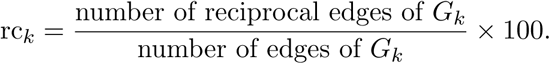

We plot the curves *k* ↦ rc*_k_* for all connectomes and their corresponding controls in Figure 2D.

The simplicial structure of *G* can be used to compute (weighted) averages of any node based property as follows. Let *N* be the number of nodes of *G* and 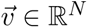 be a vector given by a node based property, e.g., node degree or reliability. For any dimension *k* and any possible position 0 *≤ pos ≤ k* within a *k*-simplex, we denote by mean(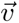, *k*, *pos*) the mean of the values of 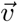 of the nodes in position *pos* of *k*-simplices, counted with multiplicity. More precisely, if *s*^1^*, …, s^t^* is a list of all *k*-simplices, let 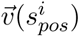 be the value of 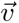 of the node in position *pos* of *s^i^*, then we define mean(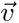*, k, pos*) as the mean

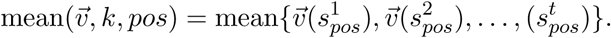

We denote by mean(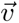*, k, all*) the weighted mean of 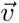 on the nodes that are in *k*-simplices (counted with multiplicity) regardless of position, i.e.,

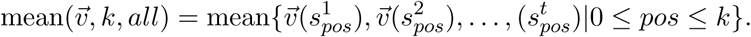

We plot the curves *k* ↦ mean(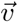*, k, all*) in Figures 3B, 6C, 4A/C for different structural and activity properties of the nodes. In the last three cases, we further plot *k* ↦ mean(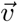*, k, pos*) for *pos* = 0 or *pos* = *k* i.e., the source respectively target neurons of a simplex.

#### 5.2.6 Neighborhood complexity

Let *G* be a network and *v* be a node in *G*. We call the *neighborhood of v* the subgraph of *G* on all the nodes in *G* that are connected to *v* and the edges between them. We denote this subgraph by *N_v_* and we call *v* its *center*. The *size of N_v_* is the number of nodes in it. In the absence of reciprocal connections, the size of *N_v_* is the total degree of *v* plus 1. However, in general we have that

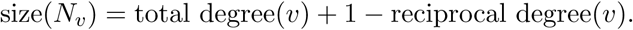

The complexity of *N_v_* is determined by how similar or different it is from a random subgraph of a similar size. In order to quantify this, we generate a configuration model control of *G*. We denote this control *Ĝ* and by *N̂_v_* the neighborhood of *v* in *Ĝ*. Note that by construction *N_v_* and *N̂_v_* are of similar size, particularly for large sparse graphs. Thus, we define the *neighborhood complexity of N_v_* as the distance between *N_v_* and *N̂_v_*. See Figure S5 for an explicit example. We measure this distance using simplex counts and define

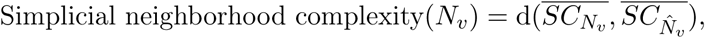

where 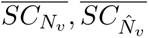 denote the simplex counts of *N_v_* and *N̂_v_* respectively linearly normalized to jointly lie between 0 and 1, and d denotes Euclidean distance. To test the robustness of our metric, we consider additionally a distance given by a metric that only takes into account pairwise interactions and define a degree based complexity metric as

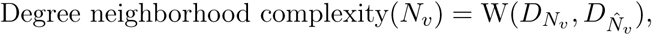

where 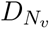 and 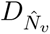 are the total degree distributions of *N_v_* and *N_v_* respectively and *W* is the Wasserstein distance between them, a classical distance between probability distributions^102^. These two values are strongly correlated to each other (Fig. S6D).

#### 5.2.7 Reliability

We measured the reliability of spike trains to repeated presentations of natural movies in the MICrONS dataset, with the *Oracle score* reported by MICrONS^49^. The score was computed as the jackknife mean of correlations between the leave-one-out mean across repeated stimuli with the remaining trial. The sparsity of the activity data set required us to pool Oracle scores across sessions for some analyses. Since the baseline Oracle scores have large variability across sessions (Fig. S9A), we first z-scored the Oracle scores in each session and then average them across sessions.

To measure the spike timing reliability of a neuron in the BBP data set, we used the notion of *Gaussian kernel reliability* ^66^ across repetitions of the same experiment. The repetitions were obtained from repeated simulations with different simulator seeds. Specifically, for a set of spike trains of a given neuron obtained from repetitions of the same experiment, we first convolved the spike trains with a Gaussian kernel with *σ* = 10 ms. Since reliability is known to be confounded by the firing rate of a neuron^103^, we removed this dependency by *mean-centering* the data (see Fig. S8), i.e., by subtracting the mean of the convolved signal. Then, for each neuron we computed the cosine similarity between all pairs of repetitions of mean-centered signals and averaged the resulting values.

#### 5.2.8 Connectome manipulation pipeline

We employed the recently developed `Connectome-Manipulator` Python framework ^67^, which allows rapid manipulations of the connectivity of circuits in the SONATA format ^68^, which we will denote by *sonata-connectomes*. The framework supports the creation of new sonata-connectomes (referred to as wiring), transplantation of connectivity features from other sonata-connectomes, and manipulations of existing sonata-connectomes (referred to as rewiring) on the level of connections and synapses. To create a manipulated sonata-connectome we require a connection probability model defining the connectivity together with parameter models specifying the physiological properties of the generated connectivity. We provide details for these in the following subsections. The adjacency matrices of all manipulated sonata-connectomes are available on Zenodo ^96^.

##### Connection probability models

There were four categories of manipulations done on the baseline connectome: *remove reciprocity, increase reciprocity, reduce complexity, enhance complexity*. These are listed in Table 2 and we describe them explicitly here. All manipulations were restricted to connections between excitatory neurons in the central hexagonal column of the circuit.

**Table 2:** Types of connectome manipulations.

| Label | Modification category | Description |
| --- | --- | --- |
| *Order 1* | Reduce complexity | Uniform probability of connection |
| *Order 2* | Reduce complexity | Distance-dependent probability of connection |
| *Order 3* | Reduce complexity | Bipolar distance-dependent probability of connection |
| *Order 4* | Reduce complexity | Offset-dependent probability of connection |
| *Order 5* | Reduce complexity | Position- and offset-dependent probability of connection |
| *RC - 1* | Remove reciprocity | Remove reciprocal connections in maximal dimension $n \geq 5$ |
| *RC - 2* | Remove reciprocity | Remove reciprocal connections in maximal dimension $n \geq 5$ and 50% at $n = 4$ |
| *RC - 3* | Remove reciprocity | Remove all reciprocal connections in dimension $n \geq 4$ |
| *RC - 4* | Remove reciprocity | Remove all reciprocal connections |
| *RC \* -K* | Remove reciprocity | Remove the same number of reciprocal connections as in *RC - K* at random, for $1 \leq K \leq 3$ |
| *RC + K* | Add reciprocity | Increase reciprocal connections on maximal simplices by a factor of $K + 1$, for $1 \leq K \leq 4$ |
| *RC + 5* | Add reciprocity | Increase reciprocal connections on maximal simplices by a factor of 8 |
| *RC \* +K* | Add reciprocity | Add the same number of connections as in *RC + K* at random, for $1 \leq K \leq 5$ |
| *Nk* | Enhance complexity | Rewiring of $Nk$ edges to increase edge participation, for $N=100, 200, 300, 400, 500, 670$ |

In case of manipulations that *reduce complexity*, labeled *Order M*, we globally rewired the connectome across the complexity spectrum based on five simplified stochastic connection probability models ^104,67^ which had been fit against the baseline connectome beforehand, using the `Connectome-Manipulator` framework. The total numbers of connections in these simplified connectomes were closely matched to the number of connections in the baseline connectome (relative difference less than 0.1 %) using the iterative procedure described in Pokorny et al. ^67^. Self-connections (autapses) were not allowed. We used the following simplified models:

**Order 1** Uniform connection probability between all pairs of pre- and post-synaptic neurons.
**Order 2** Distance-dependent connection probability; the probability depends on the Euclidean distance between pre- and post-synaptic neurons (conceptually equivalent to the *Distance model* of Table 1, but with the technical difference that it was modelled by the sum of a proximal and distal exponential function^67^). For fitting the model, we used a distance binning of 50 *µ*m.
**Order 3** Bipolar distance-dependent connection probability; same as *Order* 2 (including the binning parameters) but with a distinction between pre-synaptic neurons above or below post-synaptic neurons (in axial direction perpendicular to the cortical layers).
**Order 4** Offset-dependent connection probability; the probability depends on the axial and radial offsets between pre- and post-synaptic neurons (in a cylindrical coordinate system with axial direction perpendicular to the cortical layers). For fitting the model, we used an offset binning of 50 *µ*m.
**Order 5** Position- and offset-dependent connection probability; same as *Order* 4 but with additional dependence on the absolute axial position (i.e., cortical depth) of the pre-synaptic neuron. For fitting the model, we used an offset binning of 100 *µ*m, and a position binning of 400 *µ*m.

The other three categories of manipulations were based on adjacency matrices generated beforehand which we employed as deterministic connection probability models, i.e., containing only probability values of zeros and ones as determined by the matrix. We describe how these adjacency matrices where generated.

On the manipulations that *reduce reciprocity*, labeled *RC − K*, for selected reciprocal connections we removed one of the two directed edges at random. The reciprocal connections were selected from the spine^*^ of maximal *n*-simplices as follows:

**RC** *−* **1** Reciprocal connections on the spine of maximal *n*-simplices for *n ≥* 5.
**RC** *−* **2** Reciprocal connections on the spine of maximal *n*-simplices for *n ≥* 5 and half (chosen at random) of the reciprocal connections that are on the spine of maximal *n*-simplices for *n* = 4 that are not in the spine of a higher dimensional simplex.
**RC** *−* **3** Reciprocal connections on the spine of maximal *n*-simplices for *n ≥* 4.
**RC** *−* **4** All reciprocal connections.

To control for the effect of the location of reciprocal connections we generated manipulations, denoted *RC ∗ −K* for 1 *≤ K ≤* 3. In each manipulation *RC ∗ −K* we removed the same number of reciprocal connections as in *RC ∗ −K*, but we did so at random rather than from maximal simplices of top dimension. Note in particular, that there is no *RC ∗ −*4, since it is not possible to remove all reciprocal connections at random.

On the manipulations that *increase reciprocity*, labeled *RC* + *K*, we added reciprocal connections to existing edges such that the percentage of reciprocal connections is approximately multiplied by a chosen factor *F_K_* across all maximal simplices. To do this, let *G* be the original connectome, let *G_k_* be the subgraph on edges that are on simplices of dimension *k* (see Sub-section 5.2.5) and let *nrc_k_* be the number of reciprocal connections of *G_k_*. For all dimensions *k ≥* 2, we selected at random *F ∗ nrc_k_* non-reciprocal edges of *G_k_* that we turned into reciprocal connections. We did this for *F_K_* = 2, 3, 4, 5 and = 8. To control for the effect of increased connectivity density we generated manipulations, denoted *RC ∗* +*K* for all *K*, where we added the same number of connections as in *RC* + *K* at random in the excitatory subnetwork.

Finally, on the manipulations that *enhance complexity*, we use a custom algorithm presented by Reimann et al. ^91^ which exchanges edges in order to maximize the number of simplices an edge participates in. Briefly, for each edge in the network, we computed its *maximal dimension of edge participation*, which is the maximum dimension of simplices it participates in. For each node in the network and we computed its *n-node participation* for all dimensions *n*, which is the number of *n*-simplices the node participates in. Then we removed *M* edges at random with maximal dimension edge participation smaller or equal to 3 (the middle dimension for this connectome) and added *M* new edges at random, where the probability of an edge *a → b* to be added was proportional to the participation of *a* and *b* in simplices above dimension 3. We performed these manipulations for *M* = 100*k,* 200*k,* 300*k,* 400*k,* 500*k,* and 670*k* and labelled these manipulations as *M*. Note that 670*k* is nearly 10% of all the connections.

##### Generating sonata-connectomes

When rewiring connectivity based on a given model of connection probability, existing connections may be deleted, but also new ones created which requires parameter models specifying the physiological properties of new connections. Following the pipeline of Pokorny et al. ^67^, we first fitted a “connection properties model” against the baseline sonata-connectome. This model captured parameter distributions from which physiological property values were then drawn during rewiring, such as: conductance (gamma), decay time (truncated normal), depression time (gamma), facilitation time (gamma), utilization of synaptic efficacy (truncated normal), and number of synapses per connection (discrete). All these parameter distributions were fitted in a pathway-specific way, i.e., separately for all 13 *×* 13 pairs of pre- and post-synaptic morphological types of excitatory neurons. Second, for assigning axonal delays, we fitted a “linear distance-dependent delay model” against the baseline sonata-connectome (with 50 *µ*m distance binning) where the mean synaptic delay linearly depends on the Euclidean distance between the soma of a pre-synaptic neuron and the synapse location on the post-synaptic dendrite. Finally, in case rewired connections were to be established that already existed in the baseline sonata-connectome, there were two options: keeping them or regenerating them. For the manipulations that reduce complexity (*Order M*), we re-generated all connections whereas for all other manipulations we kept existing connections as they were, i.e., we only deleted unused ones and generated new ones that did not exist before. In all cases, we used the option for reusing existing synapse positions, so that new synapses were placed on the post-synaptic dendrites at positions randomly sampled from existing synapses in the baseline sonata-connectome.

#### 5.2.9 Coupling coefficient

Following the ideas of Okun et al. ^61^, we calculated coupling coefficient of a neuron *i* as the Pearson correlation of its activity with the activity of the rest of the circuit and denoted it *c_i_*. More precisely,

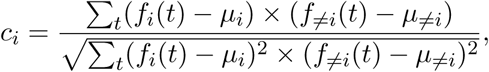

where *f_i_* is either the neuron’s deconvolved ΔF/F for MICrONS, or its binned (with 5 ms bin size) spike train for BBP, *µ_i_* is *f_i_*’*s* mean, *f*_≠*i*_(*t*) is the activity of the rest of the circuit i.e.,

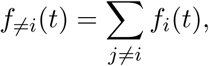

and *µ*_≠_*_i_* is its mean. Note that when the number of neurons is large, *f*_≠*i*_(*t*) is approximately the summed activity of all neurons. Thus, we used this value instead for the BBP data for an efficient implementation.

Furthermore, *c_i_* values were normalized by z-scoring their differences from the average of 10 *c_i_* values from surrogate datasets. Control datasets were created by randomly shuffling spike times, i.e., keeping single cell firing rates. We choose this *per-cell* normalization as opposed to *global* normalization (in which one would z-score *c_i_* values by the mean and standard deviation of all neurons’ surrogate *c_i_*s) as the global one resulted in a multimodal distribution for the BBP data (Fig. S14B1). The source of this artifact is that *in silico* we could record even the extremely low firing rate neurons, which lead to a low, secondary peak in the distribution (Fig. S14B2). On the other hand, for the MICrONS data set, both types of normalization returned virtually the same values (Fig. S14A).

#### 5.2.10 Efficiency of a neighborhood

We define the *efficiency of a neighborhood* as the dimension of its activity normalized by the size of its active subneighborhood i.e., the number of nodes in it that fire ^11–16^. We determined the dimensionality of the activity of the neighborhood using the correlation of the spike trains of its nodes. Thus, a neighborhood of maximum efficiency has maximally uncorrelated data and encodes more information, relative to its size.

Specifically, as for reliability, we first convolved the spike trains with a Gaussian kernel with *σ* = 10 ms, to reduce the effect of noise and we call these the *spike signals*. A fraction of all the neurons (between *∼* 25% and 30%) remain silent during a simulation. For each node *v* in *G*, let *ℓ* be the number of nodes in its neighborhood *N_v_* that fire and let Corr*_v_* denote the mean centered *ℓ × ℓ* matrix given by the pairwise Pearson correlation between the spike signals (see Fig 4B left for examples). We computed the singular values of Corr*_v_* via singular value decomposition (SVD) and we denote by 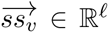 the cumulative vector of singular values normalized to lie between 0 and 1. The normalized dimension of the activity of *N_v_* is the fraction of the *ℓ* nodes required to reach 0.9 of the cumulative spectrum i.e.,

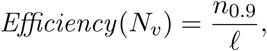

where *n*_0.9_ is the first position in 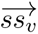 at which 0.9 is reached (see Fig 4B right).

#### 5.2.11 Classification

We used the neighborhood-based stimulus classification pipeline described by Conceição et al. ^64^ and Reimann et al. ^63^. We briefly summarize their pipeline here, which we refer to as the *single selection procedure* and refer the reader to the original papers for details on their method.

First, we computed the neighborhood metrics listed in Table S1 for the neighborhoods of all excitatory neurons within the excitatory subcircuit and called these the *selection parameters*. Second, amongst all excitatory neurons, we picked 50 neighborhood centers either: at random or those that maximize or minimize a given *selection parameter*. Third, we featurized the activity of the selected neighborhoods as a response to each stimulus and reduced it to a set of time series that we call the *feature vectors*. The featurization procedure was done in two generic ways. On one hand, following Reimann et al. ^63^, in what we call the *PCA method*, we applied principal component analysis to the binned spike trains of the union of the selected neighborhoods with a bin size of 20 ms. On the other hand, following Conceição et al. ^64^, we considered the *active subnetworks* of each neighborhood, which are the subgraphs of the neighborhood on the nodes which are active in a given time window; for each stimulus presentation we considered two time windows of 25 ms each. This reduced the activity to a set of time series of graphs (one per stimulus), which we then transformed into feature vectors by calculating their value of one of the network metrics listed in Table S1. Finally, we classified these feature vectors using a Support Vector Machine (SVM) and computed the accuracy and cross-validation error.

Note that this pipeline provides for each (selection, featurization) parameter pair a classification accuracy obtained only from the activity of 50 neighborhoods rather than the full circuit. Moreover, the sub-selection of the activity data is done purely based on the structural properties of the circuit, in particular its neighborhood structure. However, in most cases the featurization procedure uses both the structural and activity data except for the PCA method which is based in the activity values only. In particular, the number or percentage of non-zero features within the feature vectors (which are ultimately the information that enters the classifier) depend not only on the size of the union of the neighborhoods but also on the (selection, feature) pairing and (see Fig. S16).

To test the effect of the neighborhood complexity on its efficiency as a classification readout we slightly modified the pipeline above and contrasted the results with the original. The modification was simple, and affected only the second step described above in which we replaced the single selection procedure by what we call the *double selection procedure*. Specifically, instead of selecting the neighborhoods amongst all excitatory cells, we first selected those on the top 1% of reciprocal density (high complexity) or the bottom 1% (low complexity). We then selected neighborhoods from these either at random or according to a given selection parameter and followed the same steps described above. All classification results are available on Zenodo ^96^.

### 5.3 Quantification and statistical analysis

The one sample t-test was used to test if simplex counts and number of reciprocal connections of a given connectome are significantly larger/lower from the expected values on their corresponding random network controls. To compare the means of activity metrics (e.g., reliability, coupling coefficient, efficiency) across groups the Kruskal-Wallis-Test was used instead, since these distributions were not normal. In both cases, the sizes of the samples and p-values are reported in the figure captions.

To evaluate the non-symmetry of the uni-modal reliability distributions for the different manipulated connectomes, we computed their skewness using the Fisher-Pearson coefficient^105^. This parameter has a value of zero for normally distributed variables and a positive/negative value for unimodal distributions that are long tailed to the right/left.

### 5.4 Resource availability

#### 5.4.1 Lead contact

Requests for further information and resources should be directed to and will be fulfilled by the lead contact, Daniela Egas Santander.

#### 5.4.2 Materials availability

This computational study did not generate new unique materials.

#### 5.4.3 Data and code availability

We used openly available software tools listed below. To get started with recreating our analyses and understanding them better, start with the “Reliability & structure” repository. Its instructions will clarify for which parts code from other repositories are required, and which data are required.

##### Reliability & structure

Collection of tools to relate reliability, coupling coefficient and efficiency with network architecture, as well as code to create the figures for this manuscript, available under https://github.com/BlueBrain/reliability-and-structure

##### Connectome-Analysis

A library containing general functions to analyze connectomes, available under https://github.com/BlueBrain/connectome-analysis

##### Connectome-Utilities ^97^

Utilities for running topological analyses on complex networks, available under https://github.com/BlueBrain/ConnectomeUtilities

##### Connectome-Manipulator^67^

Framework for manipulations of connectomes in SONATA ^68^ format, available under https://github.com/BlueBrain/connectome-manipulator

##### TriDy/TopoSampling pipeline ^64,63^

Neighborhood-based stimulus classification pipelines, available under https://github.com/JasonPSmith/TriDy and https://github.com/BlueBrain/ topological_sampling/tree/merge_samples, respectively

##### TriDy-tools

Tools for creating new selection parameters and distributing featurization tasks for the TriDy pipeline, available under https://github.com/jlazovskis/TriDy-tools

##### assemblyfire^32^

Detection of cell assemblies, and analysis of their connectome. Only used for extracting and pre-processing the co-registered activity data of the MICrONS data set, available under https://github.com/BlueBrain/assemblyfire

All dataset are openly available as follows:

##### C. elegans connectome^47^

Available from https://www.wormatlas.org/images/NeuronType. xls and https://wormwiring.org/pages/witvliet.html respectively

##### Drosophila larva connectome^48^

Available from https://www.science.org/doi/suppl/10.1126/science.add9330/suppl_file/science.add9330_data_s1_to_s4.zip

##### MICrONS connectome^49^

Available under DOI 10.5281/zenodo.8364070 in ConnectomeUtilities^97^ format

##### BBP connectome^50^

Available under DOI 10.5281/zenodo.10599103 in ConnectomeUtilities^97^ format

##### Reliability & structure dataset^96^

Dataset containing the BBP baseline and manipulated connectomes, their simulation data, and classification results used in this manuscript, available under DOI 10.5281/zenodo.10812496

## Acknowledgements

The authors would like to thank Guillaume Tauzin for his help with Poetry related questions in relation to connalysis, Elvis Boci for his help with visualizations and Steeve Laquitaine for comments on an earlier version of this manuscript.

This study was supported by funding to the Blue Brain Project, a research center of the École polytechnique fédérale de Lausanne (EPFL), from the Swiss government’s ETH Board of the Swiss Federal Institutes of Technology.

## Author contributions

- Conceptualization: D.E.S., C.P., J.L., J.P.S., M.W.R.
- Data curation: D.E.S., C.P., A.E., J.L., J.P.S., M.W.R.
- Formal analysis: D.E.S., C.P., A.E., J.L., J.P.S., M.W.R.
- Investigation: D.E.S., C.P., J.L., J.P.S.
- Methodology: D.E.S., C.P., A.E., J.L., M.S., J.P.S., K.H., R.L., M.W.R.
- Project administration: M.W.R.
- Software: D.E.S., C.P., A.E., J.L., M.S., J.P.S., M.W.R.
- Supervision: M.W.R.
- Validation: D.E.S., C.P., A.E., J.L., J.P.S., M.W.R.
- Visualization: D.E.S., C.P., A.E.
- Writing - original draft: D.E.S., C.P., A.E., M.W.R.
- Writing - review & editing: D.E.S., C.P., A.E., J.L., M.S., J.P.S., K.H., R.L., M.W.R.

## Declaration of interests

The authors declare no competing interests but, in the spirit of transparency, note the following: Jānis Lazovskis, currently also holds a position at Printful. This affiliation was established after the majority of his contributions to this manuscript and is unrelated to the work presented here. Ran Levi also serves as the Director of MiiST-UK, Mathematical Innovation in Science and Technology, Edinburgh, Scotland, UK. This role is unrelated to this manuscript. Finally, some authors hold minimally relevant patents related to this work, which are listed below.

- Kathryn Hess: US11569978B2 and US11652603B2
- Kathryn Hess and Ran Levi: US11893471B2, US11972343B2, US20240176985A1, US20230297808A1, US20230351196A1, US11580401B2, US11663478B2, US20190378000A1, US20190378007A1
- Kathy Hess, Ran Levi and Michael Reimann: US11615

## Supplementary material

**Table S1:** Feature and selection parameters. Each parameter is marked by S, F or S+F, denoting that it is used for selection only, featurization only, or selection and featurization, respectively. We have split the parameters into three categories: *Activity*, which considers only the activity of the neighborhoods and not their network structure, *Topology*, which consider a topological metric of the active subgraphs and *Spectral*, which consider a spectral property of the active subgraphs. Detailed definitions of all network the metrics can be found in Conceição et al. ^64^.

| Abbreviation | Parameter name | Use | Type |
| --- | --- | --- | --- |
| **pca** | PCA of activity | F | Activity |
| **ns** | Neighborhood size | S+F | Activity |
| **deg** | Neuron degree | S+F | Topology |
| **ideg** | Neuron in-degree | S+F | Topology |
| **odeg** | Neuron out-degree | S+F | Topology |
| **avdeg** | Average degree | S | Topology |
| **ec** | Euler characteristic | S+F | Topology |
| **nbc** | Normalized Betti coefficient | S+F | Topology |
| **fcc** | Clustering coefficient | S+F | Topology |
| **tcc** | Transitive clustering coefficient | S+F | Topology |
| **isimplex** | i-simplex count | S | Topology |
| **icontain** | i-simplex containment | S | Topology |
| **dci** | Simplex density | S+F | Topology |
| **asg** | Adjacency spectral gap | S+F | Spectral |
| **asl** | Adjacency spectral gap (low) | S+F | Spectral |
| **asr** | Adjacency spectral radius | S+F | Spectral |
| **blsg** | Bauer Laplacian spectral gap | S+F | Spectral |
| **blsl** | Bauer Laplacian spectral gap (low) | S+F | Spectral |
| **blsr** | Bauer Laplacian spectral radius | S+F | Spectral |
| **blsRg** | Reversed Bauer Laplacian spectral gaps | S+F | Spectral |
| **blsRl** | Reversed Bauer Laplacian spectral gap (low) | S+F | Spectral |
| **blsRr** | Reversed Bauer Laplacian spectral radius | S+F | Spectral |
| **clsg** | Chung Laplacian spectral gap | S+F | Spectral |
| **clsh** | Chung Laplacian spectral gap (high) | S+F | Spectral |
| **clsr** | Chung Laplacian spectral radius | S+F | Spectral |
| **tpsg** | Transition probability spectral gap | S+F | Spectral |
| **tpsl** | Transition probability spectral gap (low) | S+F | Spectral |
| **tpsr** | Transition probability spectral radius | S+F | Spectral |
| **tpsRg** | Reversed transition probability spectral gap | S+F | Spectral |
| **tpsRl** | Reversed transition probability spectral gap (low) | S+F | Spectral |
| **tpsRr** | Reversed transition probability spectral radius | S+F | Spectral |

**Figure S1:**
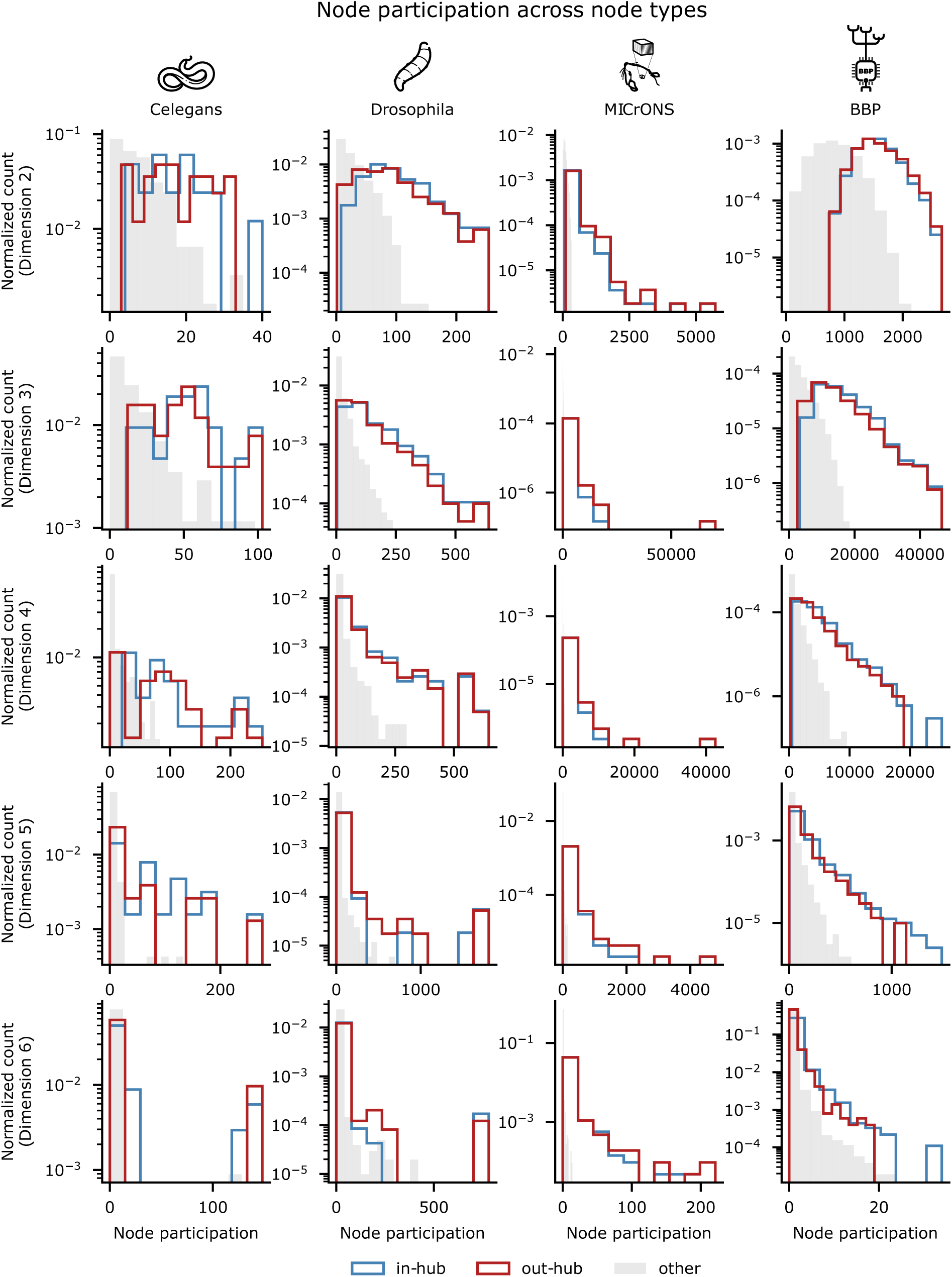
Node participation and network hubs. Normalized histograms of *k*-node participation for: all out-hubs in red, all in-hubs in blue, all other nodes in gray. In and out hubs were determined to be those in the top 10% of the in and out degree distributions. Rows correspond to the dimension of node participation 3 *≤ k ≤* 6 and columns correspond to the four connectomes.

**Figure S2:**
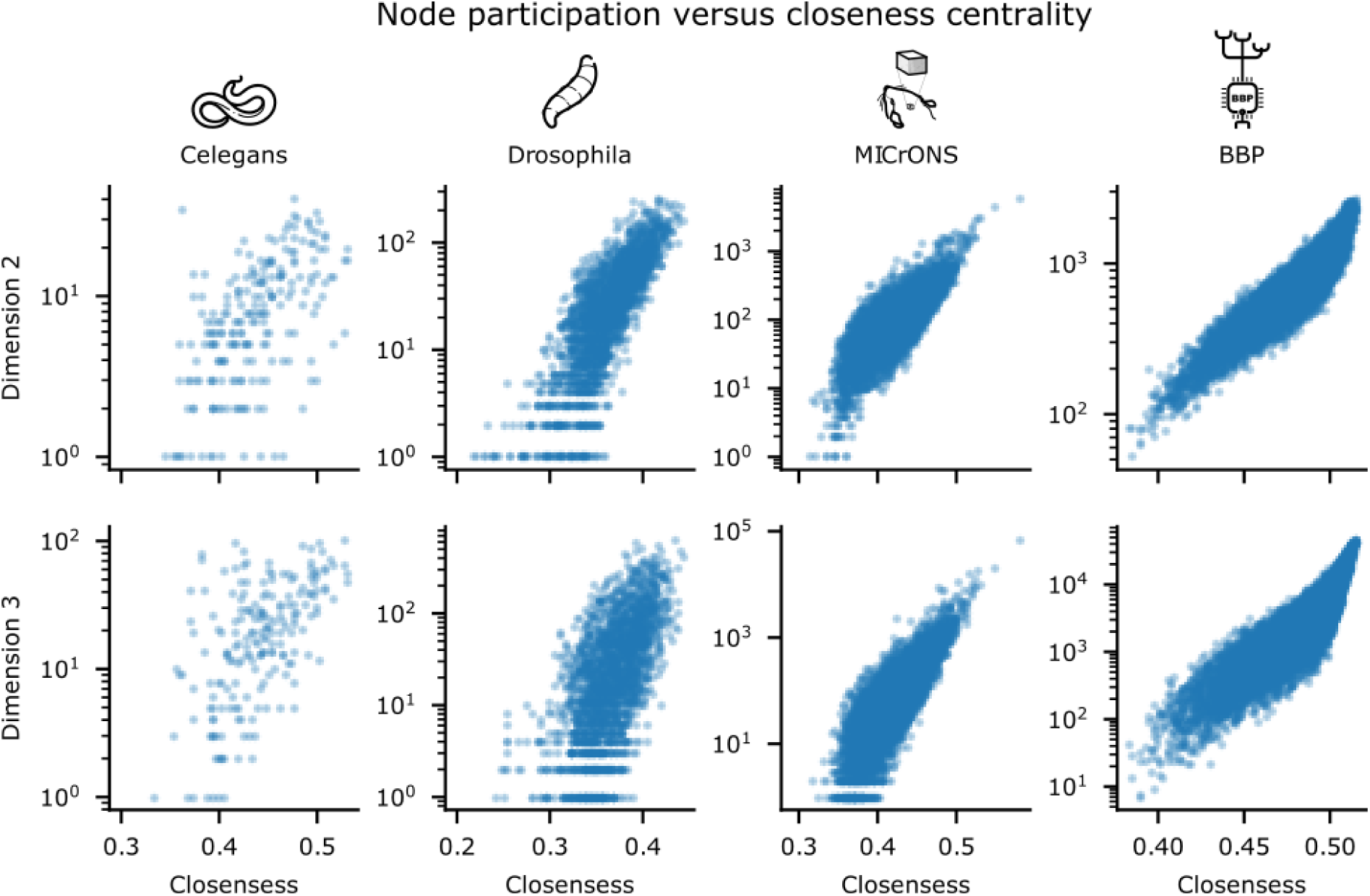
Node participation vs closeness centrality. Scatter plots of *k*-node participation (in the lower dimensions) in logarithmic scale versus closeness centrality. Rows correspond to the dimension of node participation *k* = 2, 3 and columns correspond to the four connectomes.

**Figure S3:**
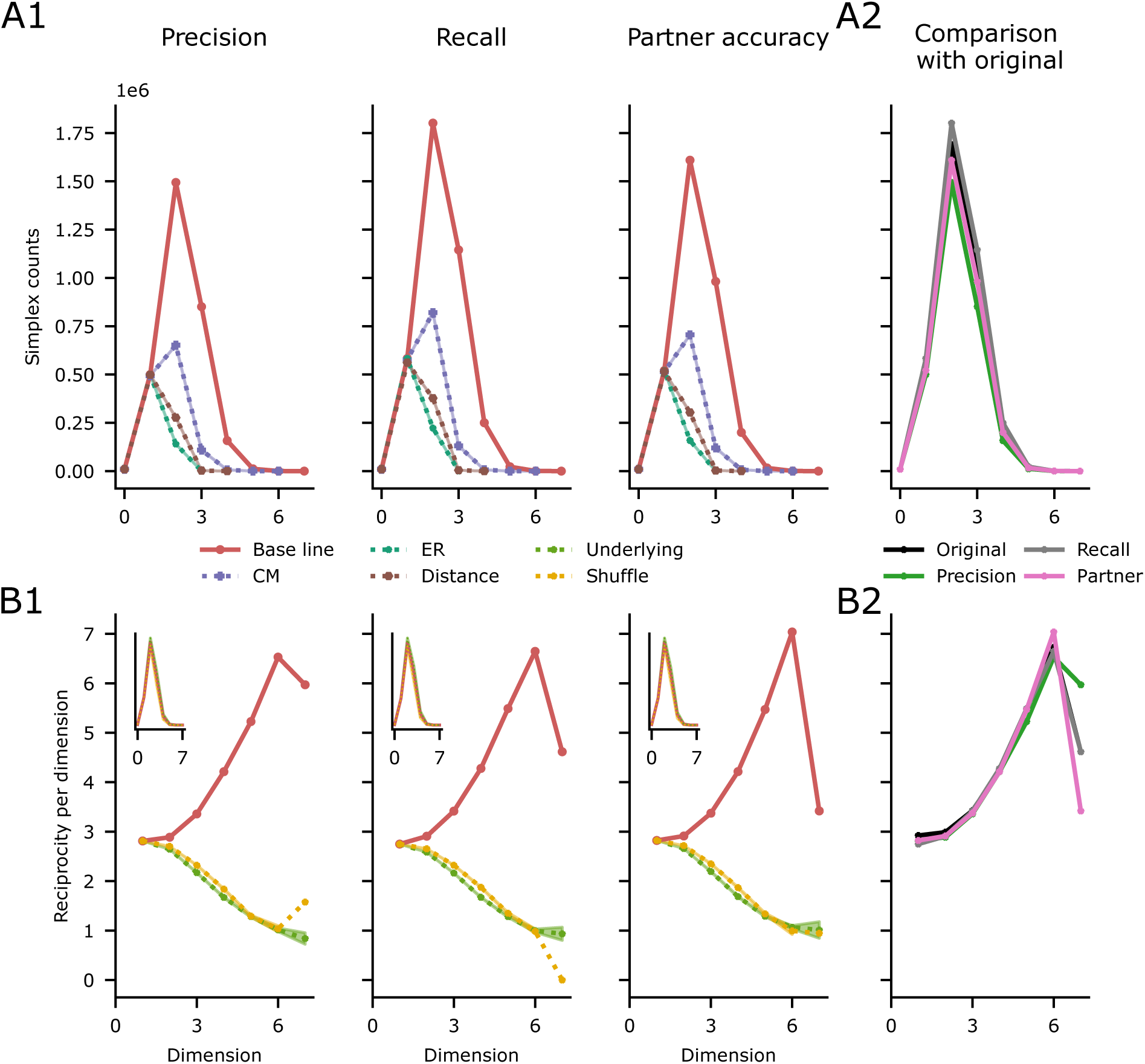
Assessing the effect of automatic detection in the MICrONS data set. Control connectomes were built to address address: precision (by removing 4% of the connections), recall (by adding 12% of connections) and partner accuracy (by shuffling 2% of the connections). Panels A1 and B1 reconstruct panels C and D of Figure 2. **A1:** Overexpression of simplex motifs with respect to 10 random controls of each type, which by design match the counts of the original network for dimensions 0 and 1. All counts in dimensions greater than one are significantly higher than the mean of the controls with p-values under 3 *×* 10*^−^*^60^ for a one sided one sample t-test. **A2:** Simplex counts for the original MICrONS connectome and the precision, recall and partner accuracy controls. **B1:** Percentage of reciprocal connections in the subgraph of simplices of each dimension contrasted with the same curve for controls where just the directionality of the connections is modified (see Methods). Inset: Simplex counts of the directional controls are close to the original ones. **B2:** Percentage of reciprocal connections on the subraphgs of simplices of the original MICrONS connectome and the precision, recall and partner accuracy controls.

**Figure S4:**
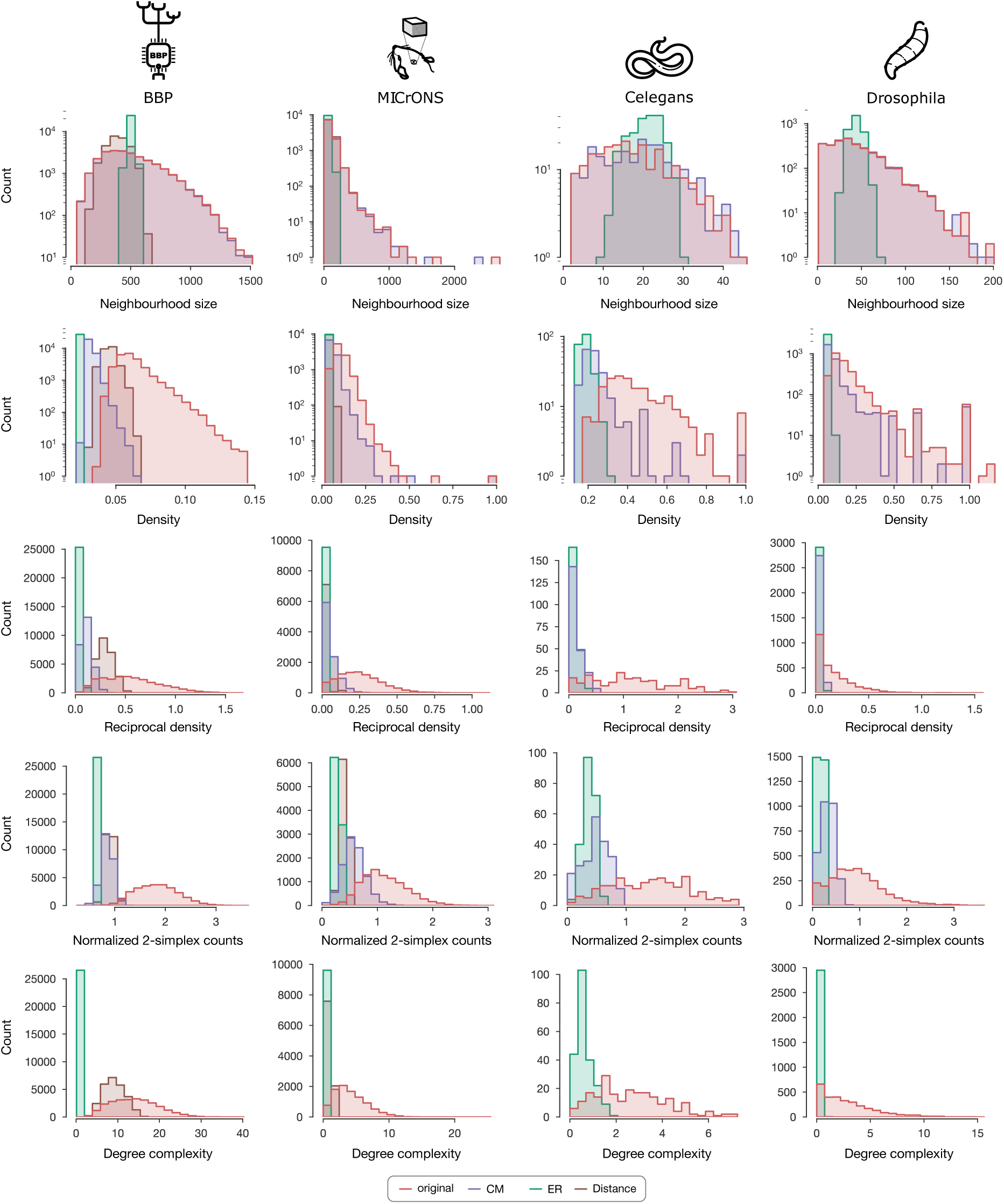
Network complexity promotes diversity of neighborhoods. Distribution of network metrics across neighborhoods for the original connectome and *ER*, *CM* and distance controls (when available). Row-by-row: Neighborhood size, density, reciprocal density, normalized 2-simplex counts, and degree complexity across connectomes. Normalization of 2-simplex counts was done by 1-simplex counts.

**Figure S5:**
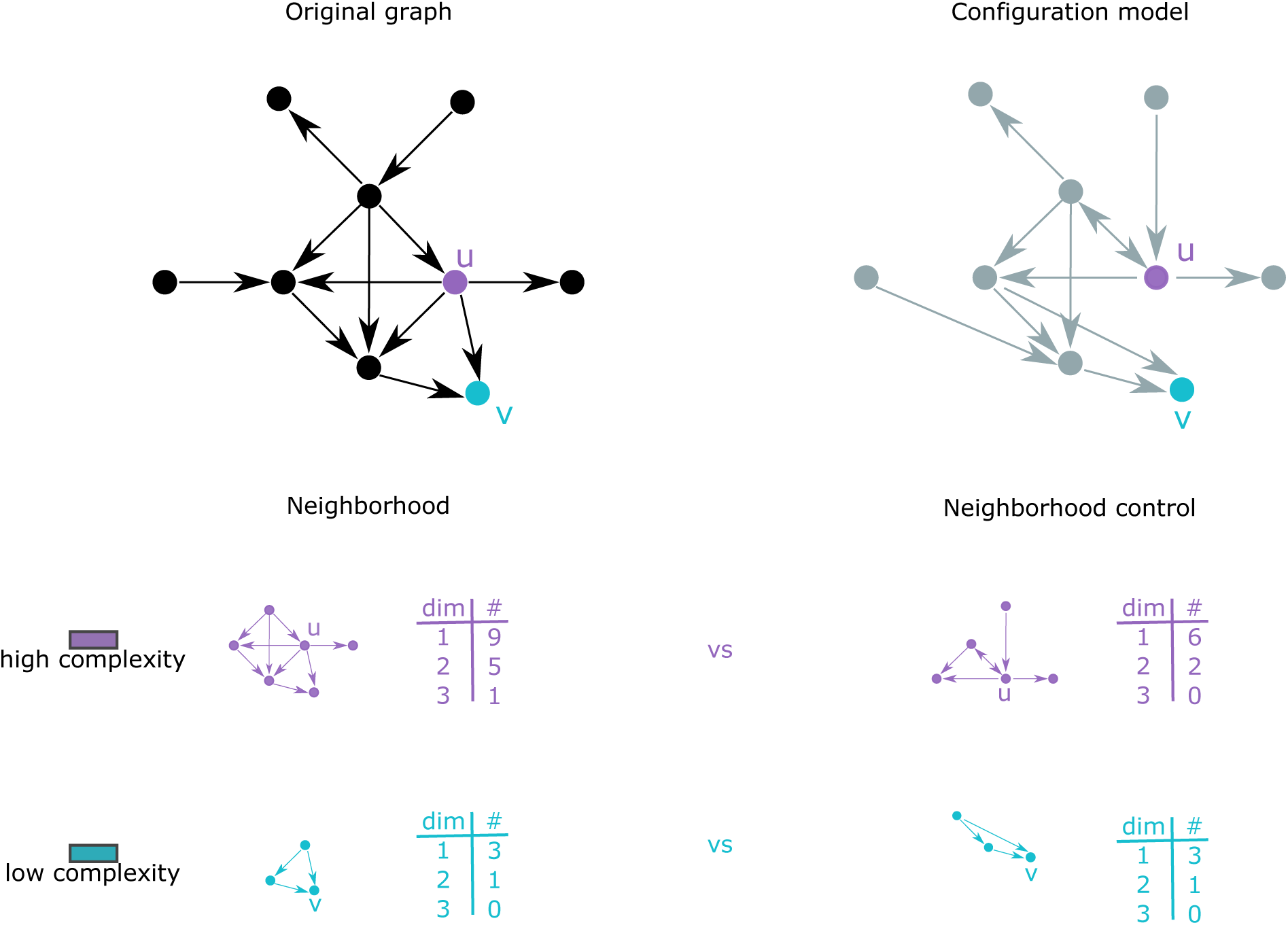
Illustrative example of simplicial/degree complexity. Top: Original base graph on the left and its corresponding control on the right. Bottom: The neighborhoods of the nodes *u* and *v* on the left and their corresponding controls on the right. The tables indicate the simplex counts in each dimension of each of the neighborhood subgraphs.

**Figure S6:**
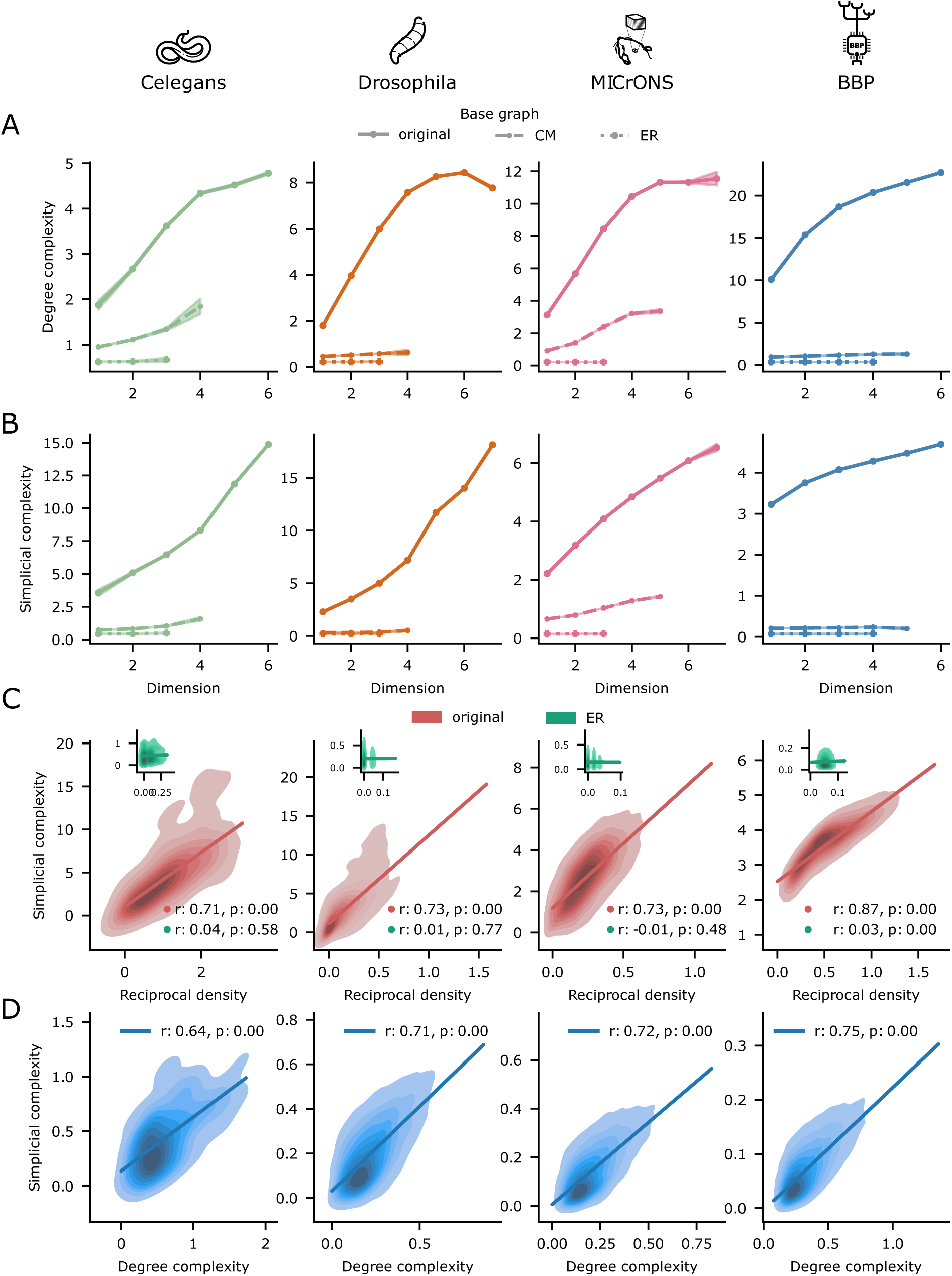
Neighborhood complexity (simplicial and degree). **A:** Average of neighborhood degree complexity on nodes that are on simplices of a given dimension(see Methods); drawn with solid, dashed and dotted lines for the original connectome a CM control and an ER control respectively. **B:** As A but with simplicial complexity instead. **C:** In BNNs reciprocal density is strongly correlated with simplicial complexity. Inset, this is not the case for ER graphs. **D:** The metrics of degree complexity and

**Figure S7:**
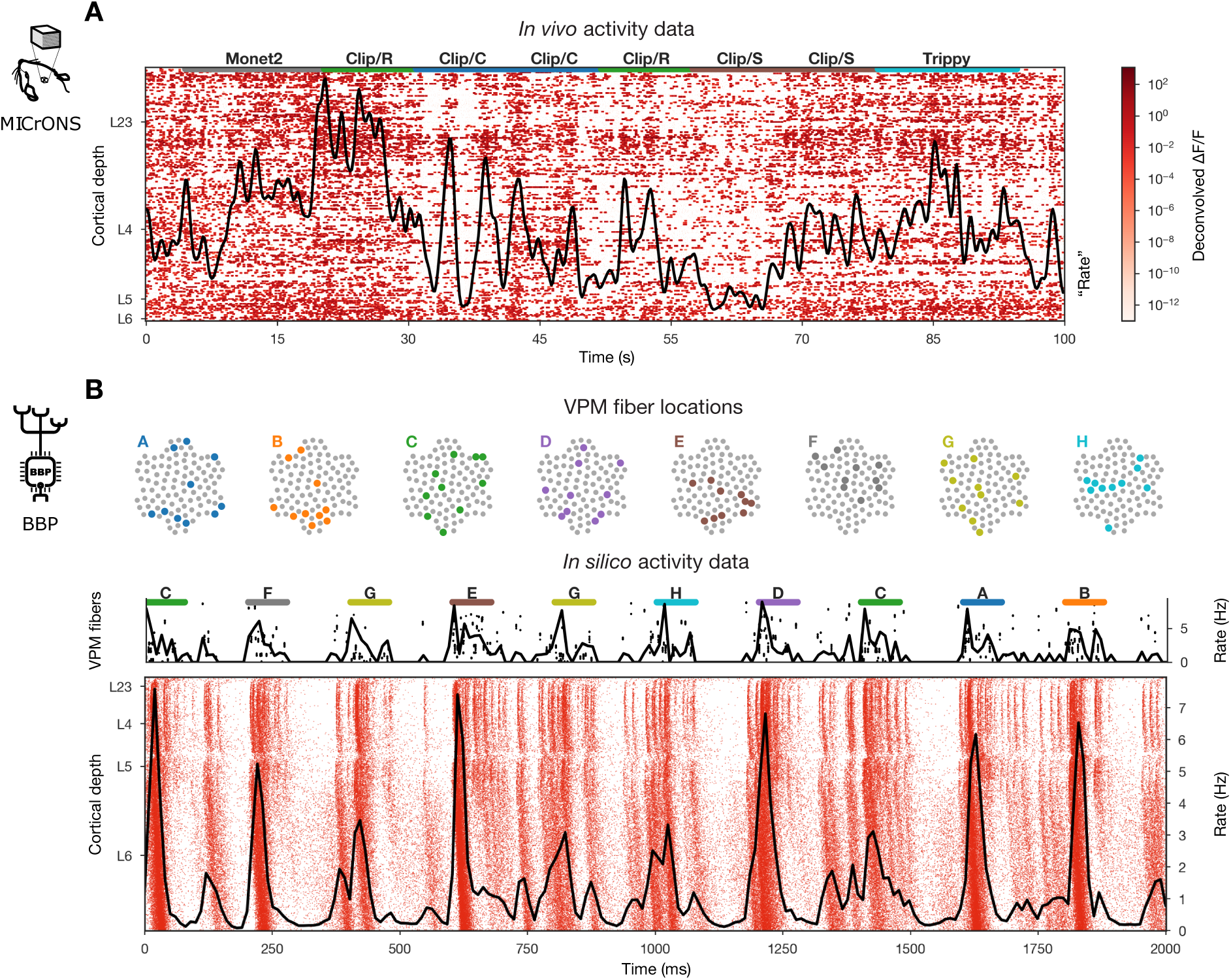
Activity datasets. Network activity and input stimuli on top for MICrONS (A) and BBP (B). **A:** MICrONS data shows deconvolved fluorescent traces, i.e., spike traces as extracted by the original authors ^49^. It is 100 seconds from scan 7 from recording session 4. **B:** VPM input fiber bundles corresponding to the eight input patterns and input raster below. Raster plot (i.e., dots represent spikes from excitatory cells) of the BBP network.

**Figure S8:**
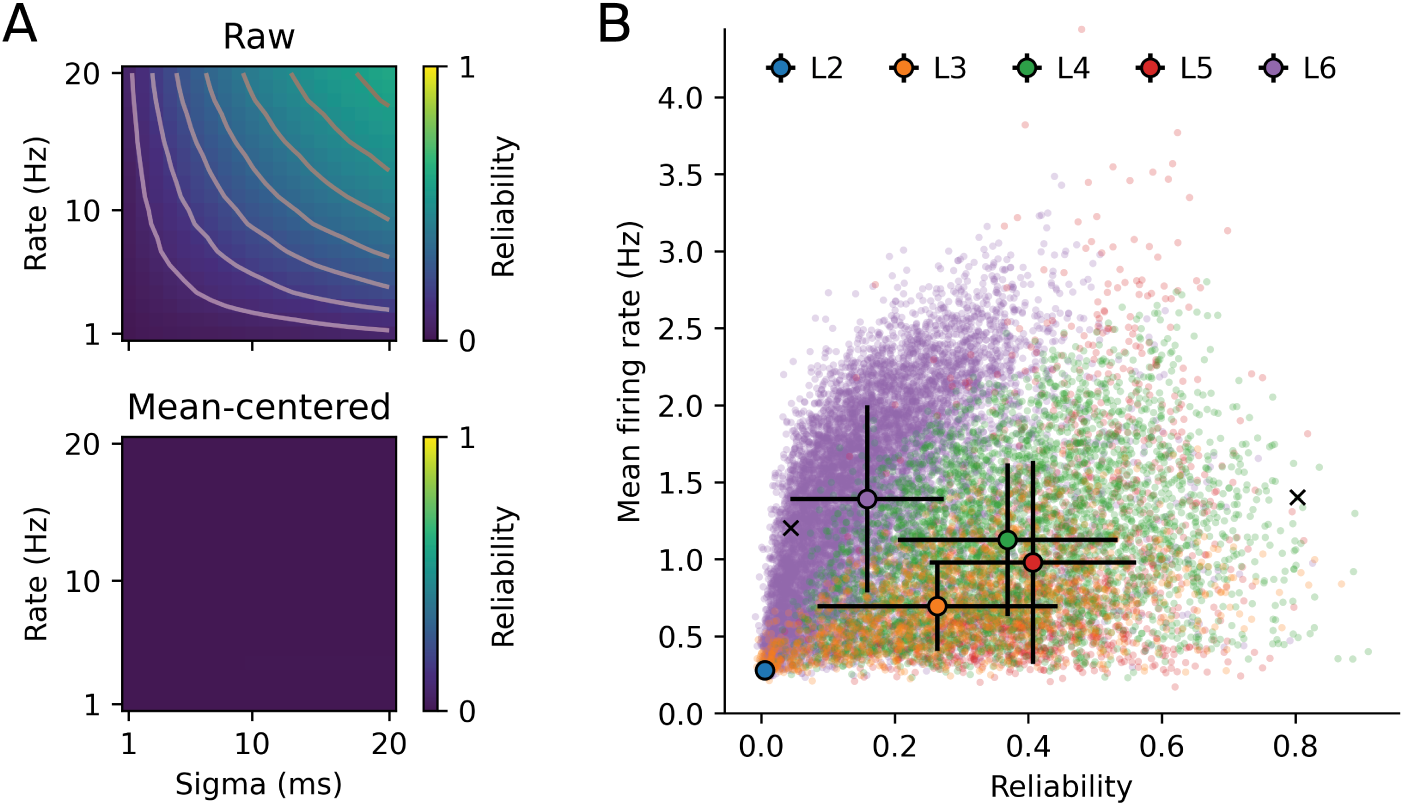
Reliability versus firing rates. **A:** Gaussian kernel reliability values of randomly generated Poisson spike trains with different firing rates and computed with different values of sigma. Without mean-centering of the data, there is a clear dependence of reliability on firing rates (upper panel) which disappears completely when mean-centering the data (lower panel). **B:** Mean-centered Gaussian kernel reliability values (*σ* = 10 ms) versus mean firing rates of all excitatory neurons in BBP data, colored by layer. Error bars denote mean *±* SD per layer. *×* markers indicate neurons with example spike trains shown in Fig. 6A.

**Figure S9:**
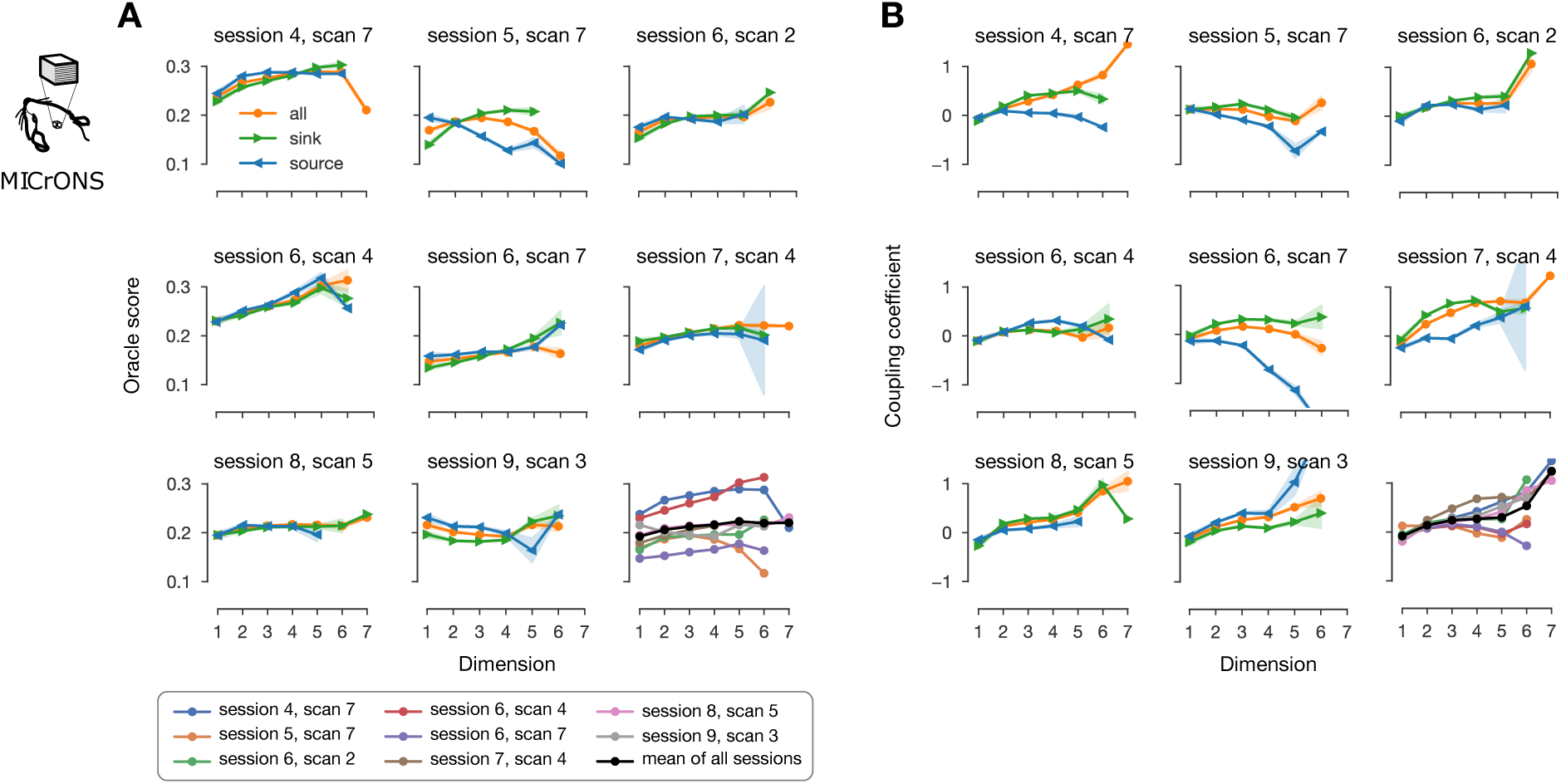
Per-scan activity metrics for MICrONS. Oracle score on panel **A** and coupling coefficient on panel **B** across simplices for the eight different recording sessions used (see Methods). The orange curves are the weighted averages of all neurons in that dimension, while the blue and green curves consider only the neurons in source or sink positions respectively. On the last subfigure, the curves for all nodes are pooled together across scans and the black line indicates the mean across sessions. (Mean across sessions is not the same as on the main figures, where the functional data was first z-scored and pooled together before running the analysis.) The same colors and legends as in A apply to B.

**Figure S10:**
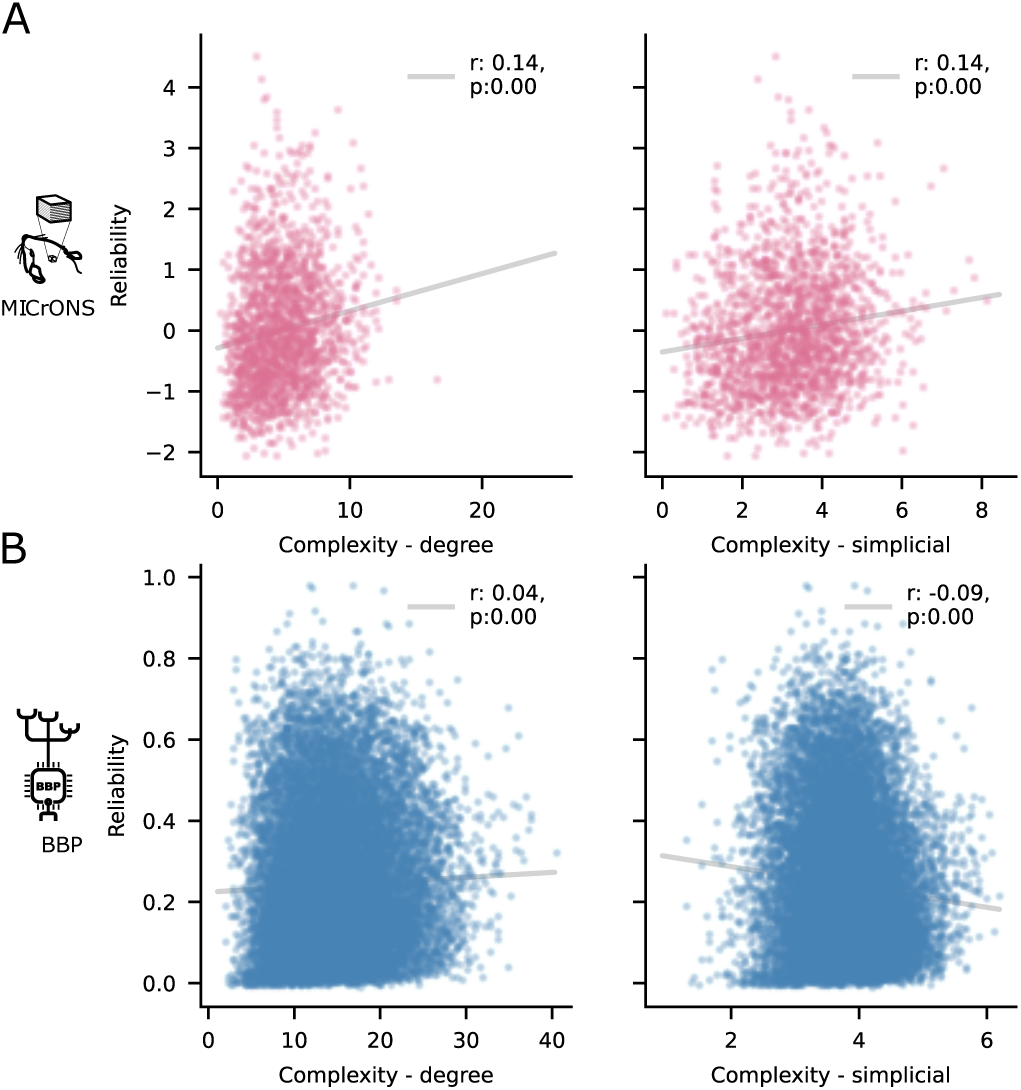
Reliability and neighborhood complexity. Negligible correlation is found between the neighborhood complexity and the reliability of its center, both for the degree and simplicial complexity metrics.

**Figure S11:**
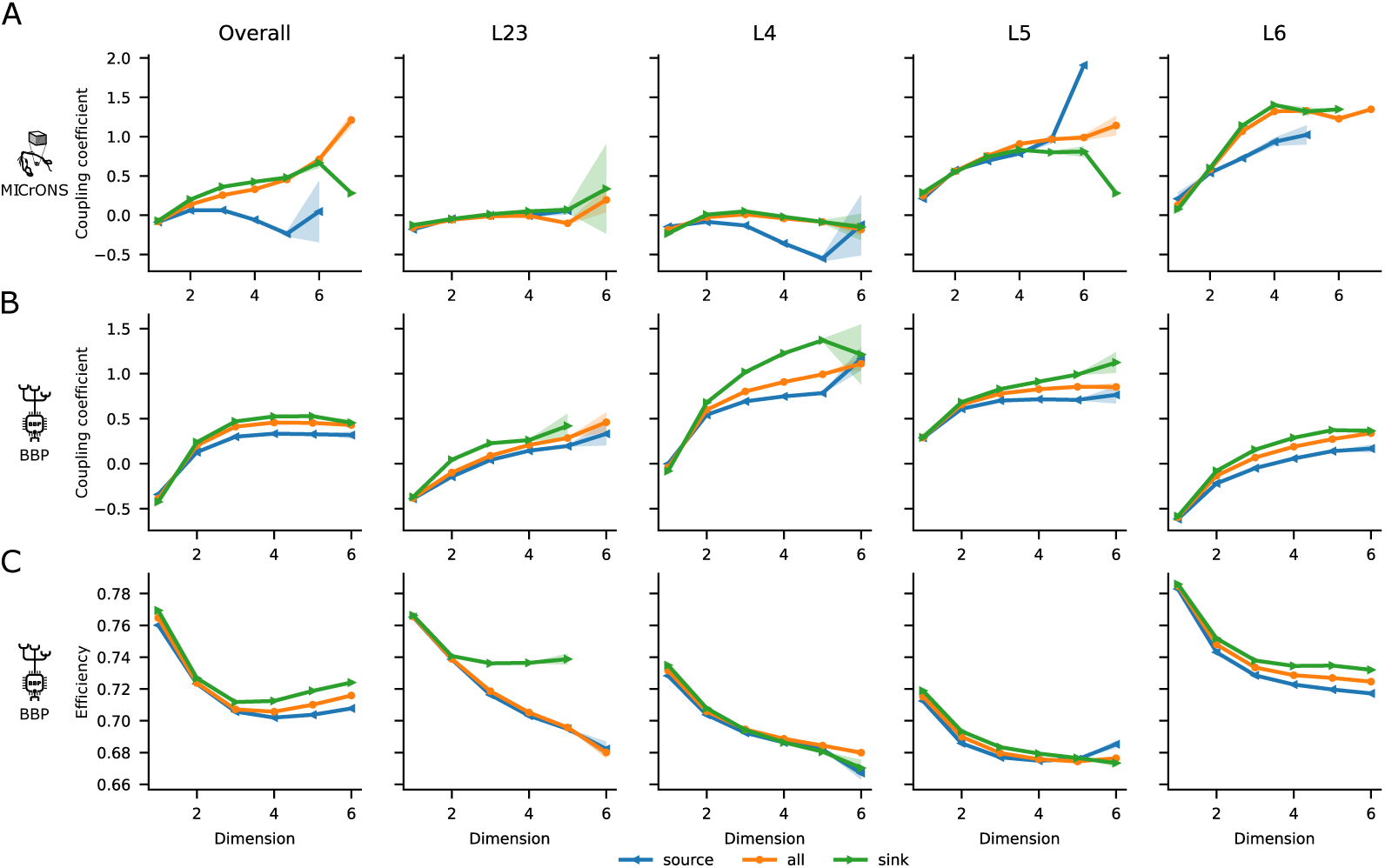
Coupling coefficient and efficiency across simplices split by layer. Coupling coefficient across simplices on panel **A** for MICrONS and **B** for BBP. On the left for all the neurons as in Figure 4A and on the right of it split across layers. **C:** As for **A** and **B** but for neighborhood efficiency (BBP data only). For all panels, the orange curves are the weighted averages of all neurons in that dimension, while the blue and green curves consider only the neurons in source or sink positions respectively.

**Figure S12:**
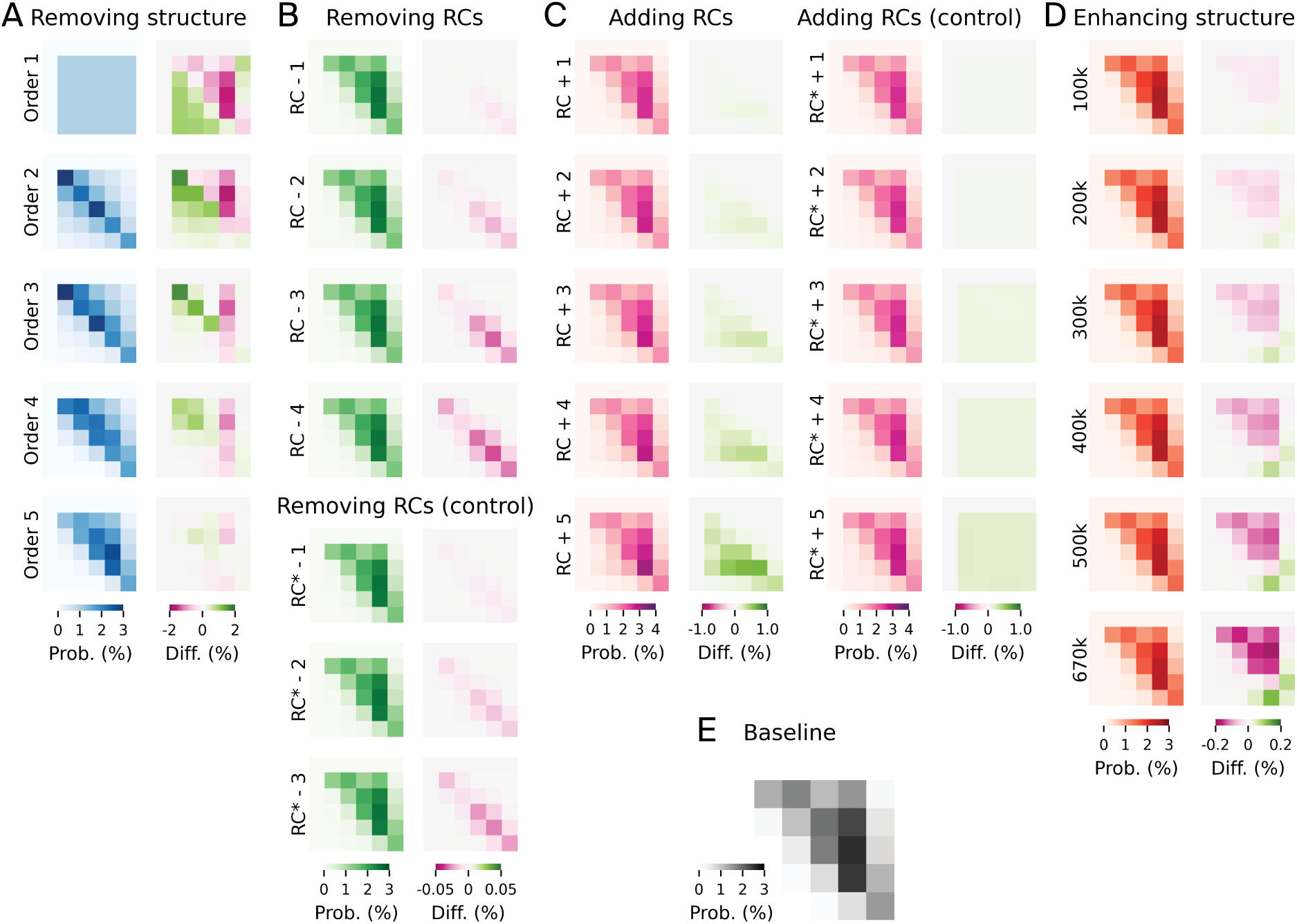
Structural validation of manipulated connectomes. For all manipulated connectomes shown in Fig. 5 and summarized in Table 2, the layer-wise connection probabilities when removing structure (**A**), removing reciprocal connections (**B**), adding reciprocal connections (**C**), and enhancing structure (**D**), and their differences to the baseline connectome (**E**) are illustrated. Asterisks indicate random controls. In the first column of each panel, the mean connection probabilities between excitatory neurons in each pre-synaptic (y axis) and post-synaptic (x axis) layer are shown. Note that layer 1 is empty. In the second column, the differences to the original connectome are highlighted. Note the different color scales at the bottom.

**Figure S13:**
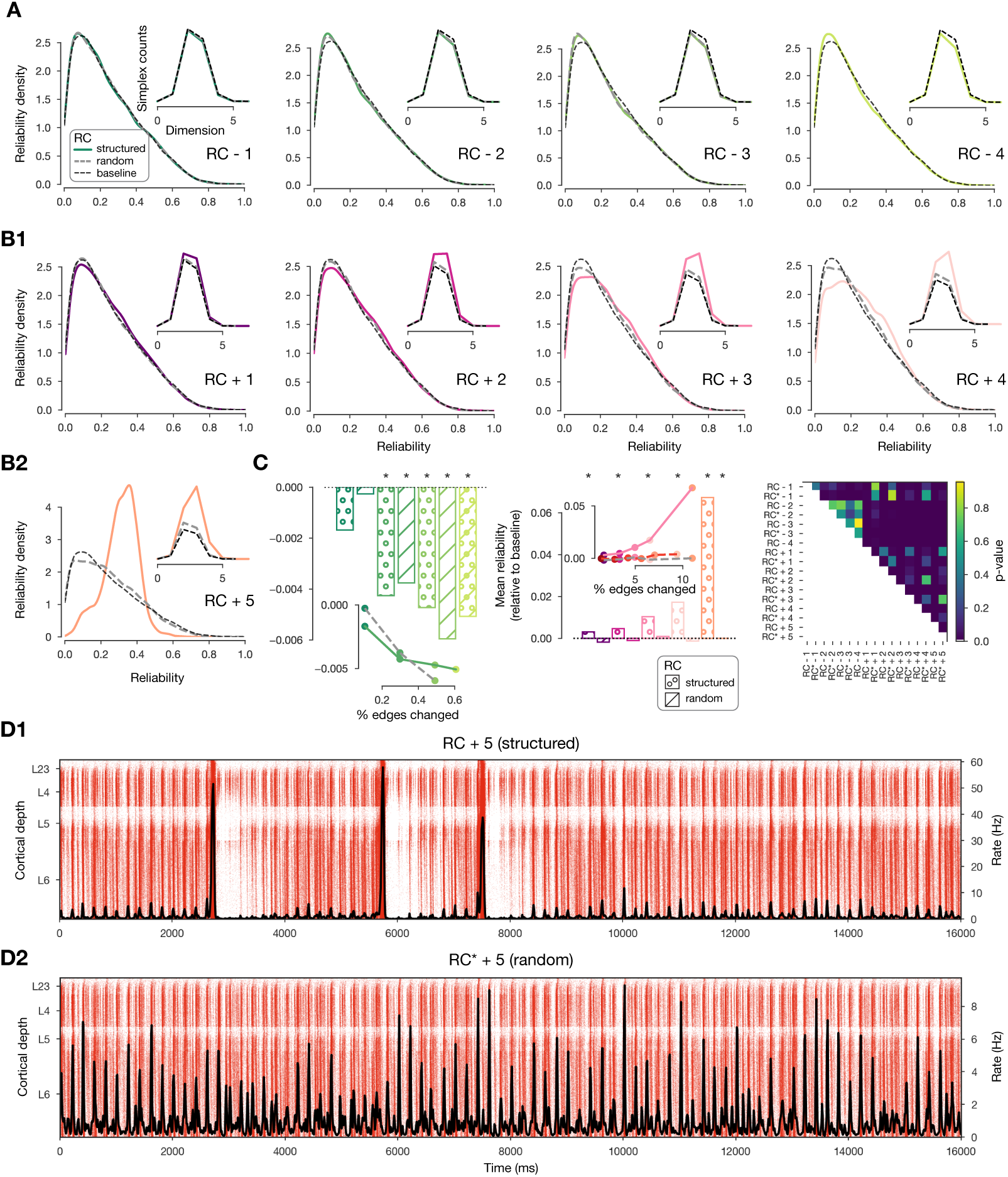
Reliability of manipulated connectomes relative to their controls. Distributions of neurons in a manipulated connectome (color), its control (dashed grey) and the baseline connectome (dashed black). Insets show corresponding simplex counts with the same color coding. Panel **A** shows manipulations where reciprocal connections are removed and their controls i.e., *RC − k* and *RC ∗−k*. See Table 2 for details. Panel **B** as A but for reciprocal connections added i.e., *RC* +*k* and *RC ∗* +*k*. **C:** Left change in mean reliability relative to the baseline connectome for reciprocal connections removed. The bars marked with a *∗* indicate when the change is statistically significant according to a KW-test. The insets show the change in mean reliability vs % of the edges that have been changed. Middle, as left but for reciprocal connections added. Right, p-values of the KW-test between all manipulated connectomes in the Figure. **D1, D2**: Raster plot of the activity in *RC* + 5 and *RC ∗* +5 respectively, showing that *RC* + 5 has transitioned to synchronous activity.

**Figure S14:**
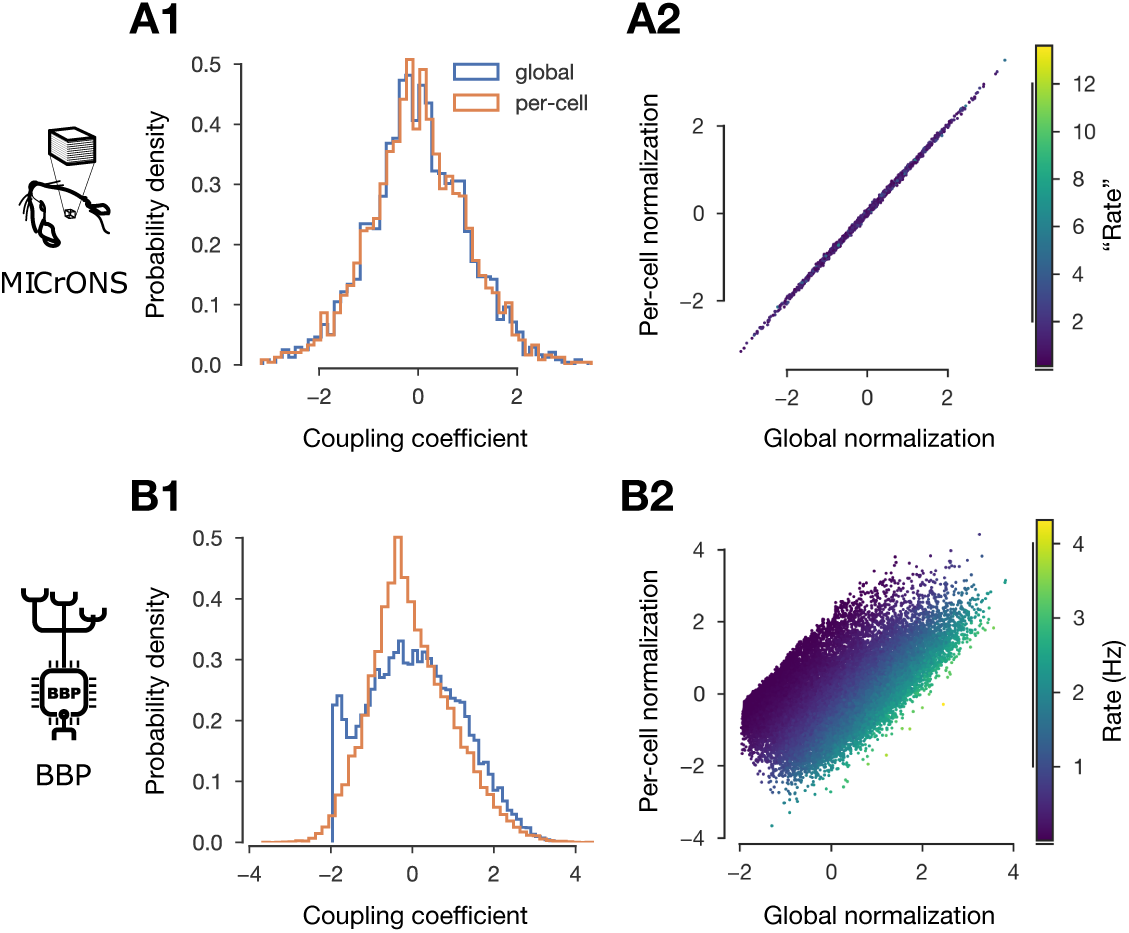
Normalization of raw coupling coefficients. **A1** and **B1:** Global vs. stricter per-cell (keeping the firing rate, see Methods) normalization of coupling coefficients for MICrONS (A1) and BBP (B1). **A2** and **B2:** Correlation between global and per-cell normalized values and their link to firing rate (especially for BBP, where the two distributions differ.).

**Figure S15:**
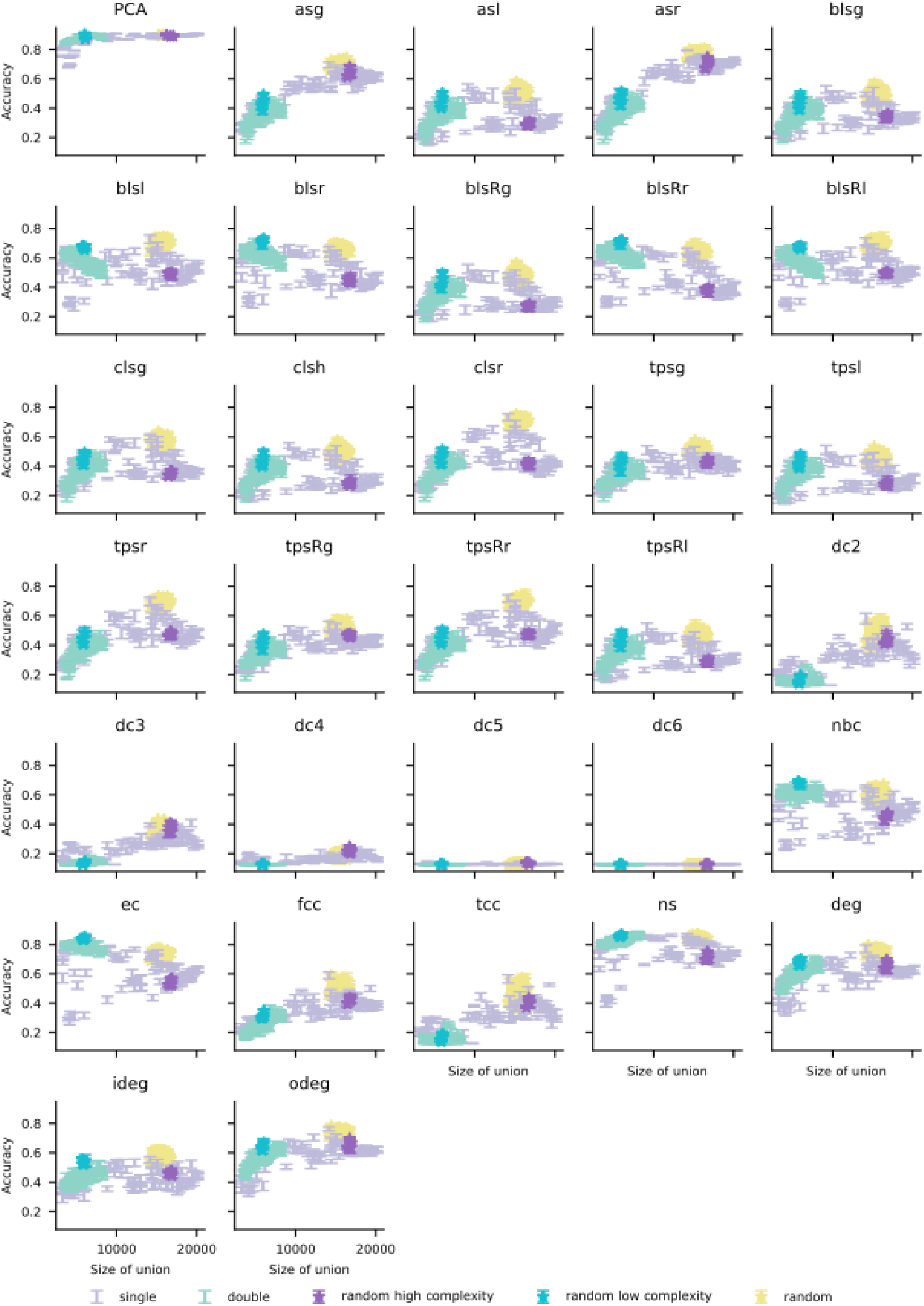
Classification accuracy for a given population size. Classification accuracy for all feature parameters. Each panel corresponds to a feature parameter, which is indicated in its title. Within each panel each bar corresponds to a selection of 50 neighborhoods selected either at random or maximizing/minimizing a given selection parameter. The x-axis of the bars corresponds to the size of the union of the 50 neighborhoods selected. Its center in the y-axis corresponds to the the accuracy for that

**Figure S16:**
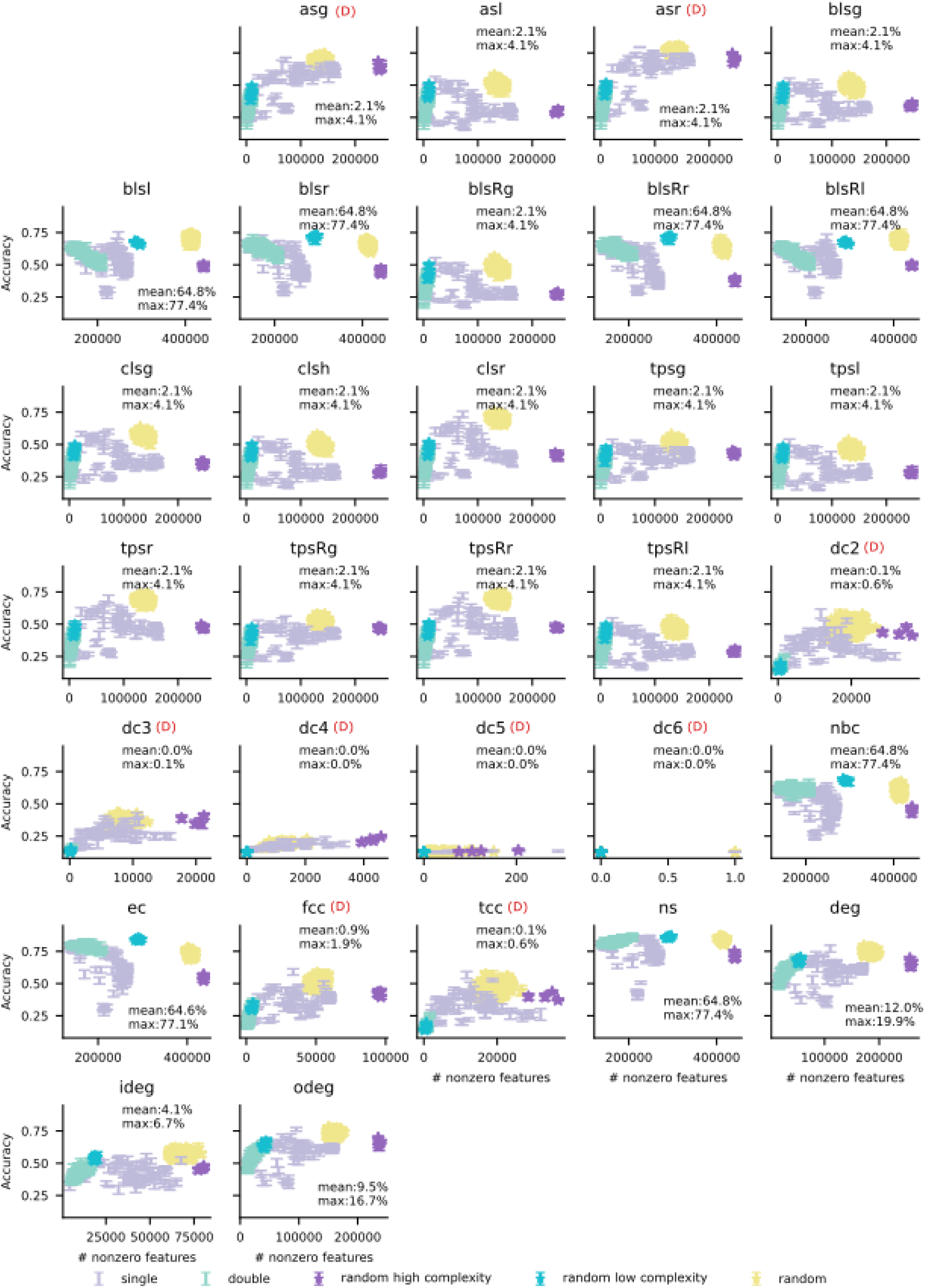
Classification accuracy versus number of nonzero features. Classification accuracy for all feature parameters as for Fig. S15 but where x-axis is given by the number of non-zero features in the featurization vector of that (selection,feature) pair. The legend indicates the mean and maximum percentage of nonzero entries in the feature vectors for the double classification procedure across all selection parameters. Degenerate featurization parameters (those for which most of the entries of their feature vectors for double selection are 0) have a (D) marking in their corresponding panels.

## Footnotes

† This is an example of the tradeoff between robustness (classification accuracy) and the efficiency (the inverse of size) presented in Figure 1B

* Recall that the spine of an *n*-simplex is the set of edges 0 *→* 1 *→* 2 *→ … → n*.

## Notes

https://doi.org/10.5281/zenodo.10812496

### Competing Interest Statement

The authors have declared no competing interest.

### Summary of Updates

Paragraphs added in the Discussion linking the current work to other areas or research.

## References

[1] Edward A Stern, Anthony E Kincaid, and Charles J Wilson. Spontaneous subthreshold membrane potential fluctuations and action potential variability of rat corticostriatal and striatal neurons in vivo. Journal of neurophysiology, 77(4):1697–1715, 1997.

[2] Michael N Shadlen and William T Newsome. The variable discharge of cortical neurons: implications for connectivity, computation, and information coding. Journal of neuroscience, 18(10):3870–3896, 1998.

[3] Majid H Mohajerani, Allen W Chan, Mostafa Mohsenvand, Jeffrey LeDue, Rui Liu, David A McVea, Jamie D Boyd, Yu Tian Wang, Mark Reimers, and Timothy H Murphy. Spontaneous cortical activity alternates between motifs defined by regional axonal projections. Nature neuroscience, 16(10):1426–1435, 2013.

[4] A Aldo Faisal, Luc PJ Selen, and Daniel M Wolpert. Noise in the nervous system. Nature reviews neuroscience, 9(4):292–303, 2008.

[5] Alfonso Renart and Christian K Machens. Variability in neural activity and behavior. Current opinion in neurobiology, 25:211–220, 2014.

[6] J Gerard G Borst. The low synaptic release probability in vivo. Trends in neurosciences, 33(6):259–266, 2010.

[7] Donald O. Hebb. The Organization of Behavior; A Neuropsychological Theory. John Wiley & Sons, Inc., New York, 1949. doi: 10.2307/1418888.

[8] Kenneth D Harris, Jozsef Csicsvari, Hajime Hirase, George Dragoi, and György Buzsáki. Organization of cell assemblies in the hippocampus. Nature, 424(6948):552–556, 2003.

[9] Vítor Lopes-dos Santos, Sidarta Ribeiro, and Adriano BL Tort. Detecting cell assemblies in large neuronal populations. Journal of neuroscience methods, 220(2):149–166, 2013.

[10] Juan A Gallego, Matthew G Perich, Lee E Miller, and Sara A Solla. Neural manifolds for the control of movement. Neuron, 94(5):978–984, 2017.

[11] John K Chapin and Miguel AL Nicolelis. Principal component analysis of neuronal ensemble activity reveals multidimensional somatosensory representations. Journal of neuroscience methods, 94(1):121–140, 1999.

[12] Eero P Simoncelli and Bruno A Olshausen. Natural image statistics and neural representation. Annual review of neuroscience, 24(1):1193–1216, 2001.

[13] Christian K Machens, Ranulfo Romo, and Carlos D Brody. Functional, but not anatomical, separation of “what” and “when” in prefrontal cortex. Journal of Neuroscience, 30(1):350–360, 2010.

[14] John P Cunningham and Byron M Yu. Dimensionality reduction for large-scale neural recordings. Nature neuroscience, 17(11):1500–1509, 2014.

[15] Peiran Gao, Eric Trautmann, Byron Yu, Gopal Santhanam, Stephen Ryu, Krishna Shenoy, and Surya Ganguli. A theory of multineuronal dimensionality, dynamics and measurement. BioRxiv, page 214262, 2017.

[16] Carsen Stringer, Marius Pachitariu, Nicholas Steinmetz, Matteo Carandini, and Kenneth D Harris. High-dimensional geometry of population responses in visual cortex. Nature, 571 (7765):361–365, 2019.

[17] Fred Attneave. Some informational aspects of visual perception. Psychological review, 61 (3):183, 1954.

[18] Horace B Barlow et al. Possible principles underlying the transformation of sensory messages. Sensory communication, 1(01):217–233, 1961.

[19] J. J. Hopfield. Neural networks and physical systems with emergent collective computational abilities. PNAS, 79(8):2554–2558, 1982. doi: 10.1073/pnas.79.8.2554.

[20] Dmitry Krotov and John J. Hopfield. Dense associative memory for pattern recognition. Advances in Neural Information Processing Systems, 29:1172–1180, 2016.

[21] Elizabeth Gardner. The space of interactions in neural network models. Journal of physics A: Mathematical and general, 21(1):257, 1988.

[22] Nils Bertschinger, Thomas Natschläger, and Robert Legenstein. At the edge of chaos: Realtime computations and self-organized criticality in recurrent neural networks. Advances in neural information processing systems, 17, 2004.

[23] Robert Legenstein and Wolfgang Maass. Edge of chaos and prediction of computational performance for neural circuit models. Neural networks, 20(3):323–334, 2007.

[24] Bosiljka Tadić and Roderick Melnik. Self-organised critical dynamics as a key to fundamental features of complexity in physical, biological, and social networks. Dynamics, 1(2): 181–197, 2021.

[25] Dietmar Plenz, Tiago L Ribeiro, Stephanie R Miller, Patrick A Kells, Ali Vakili, and Elliott L Capek. Self-organized criticality in the brain. Frontiers in Physics, 9:639389, 2021.

[26] Ashok Litwin-Kumar and Brent Doiron. Slow dynamics and high variability in balanced cortical networks with clustered connections. Nature Neuroscience, 15(11):1498–1505, November 2012. ISSN 1097-6256, 1546-1726. doi: 10.1038/nn.3220. URL https://www.nature.com/articles/nn.3220.

[27] Gustavo Deco and Etienne Hugues. Neural Network Mechanisms Underlying Stimulus Driven Variability Reduction. PLoS Computational Biology, 8(3):e1002395, March 2012. ISSN 1553-7358. doi: 10.1371/journal.pcbi.1002395. URL https://dx.plos.org/10.1371/journal.pcbi.1002395.

[28] Fereshteh Lagzi and Stefan Rotter. Dynamics of Competition between Subnetworks of Spiking Neuronal Networks in the Balanced State. PLOS ONE, 10(9):e0138947, September 2015. ISSN 1932-6203. doi: 10.1371/journal.pone.0138947. URL https://dx.plos.org/10.1371/journal.pone.0138947.

[29] Friedemann Zenke, Everton J. Agnes, and Wulfram Gerstner. Diverse synaptic plasticity mechanisms orchestrated to form and retrieve memories in spiking neural networks. Nature Communications, 6(6922), 2015. doi: 10.1038/ncomms7922.

[30] Max Nolte, Michael W Reimann, James G King, Henry Markram, and Eilif B Muller. Cortical reliability amid noise and chaos. Nature communications, 10(1):3792, 2019.

[31] Michael W Reimann, Max Nolte, Martina Scolamiero, Katharine Turner, Rodrigo Perin, Giuseppe Chindemi, Paweł Dłotko, Ran Levi, Kathryn Hess, and Henry Markram. Cliques of neurons bound into cavities provide a missing link between structure and function. Frontiers in computational neuroscience, 11:48, 2017.

[32] András Ecker, Daniela Egas Santander, Sirio Bolaños-Puchet, James B. Isbister, and Michael W. Reimann. Cortical cell assemblies and their underlying connectivity: an in silico study. PLoS Computational Biology, 20(3):e1011891, 2024. doi: 10.1371/journal.pcbi.1011891.

[33] Ann E Sizemore, Chad Giusti, Ari Kahn, Jean M Vettel, Richard F Betzel, and Danielle S Bassett. Cliques and cavities in the human connectome. Journal of computational neuroscience, 44:115–145, 2018.

[34] Bosiljka Tadić, Miroslav Andjelković, and Roderick Melnik. Functional geometry of human connectomes. Scientific reports, 9(1):12060, 2019.

[35] Ann E Sizemore, Jennifer E Phillips-Cremins, Robert Ghrist, and Danielle S Bassett. The importance of the whole: topological data analysis for the network neuroscientist. Network Neuroscience, 3(3):656–673, 2019.

[36] Miroslav Andjelković, Bosiljka Tadić, and Roderick Melnik. The topology of higher-order complexes associated with brain hubs in human connectomes. Scientific reports, 10(1): 17320, 2020.

[37] Dinghua Shi, Zhifeng Chen, Xiang Sun, Qinghua Chen, Chuang Ma, Yang Lou, and Guanrong Chen. Computing cliques and cavities in networks. Communications Physics, 4(1): 249, 2021.

[38] Jean-Gabriel Young, Giovanni Petri, Francesco Vaccarino, and Alice Patania. Construction of and efficient sampling from the simplicial configuration model. Physical Review E, 96 (3):032312, 2017.

[39] Florian Unger and Jonathan Krebs. Mcmc sampling of directed flag complexes with fixed undirected graphs. Journal of Applied and Computational Topology, pages 1–36, 2024.

[40] Ernesto Estrada and Grant J Ross. Centralities in simplicial complexes. applications to protein interaction networks. Journal of theoretical biology, 438:46–60, 2018.

[41] Emma K Towlson, Petra E Vértes, Sebastian E Ahnert, William R Schafer, and Edward T Bullmore. The rich club of the c. elegans neuronal connectome. Journal of Neuroscience, 33(15):6380–6387, 2013.

[42] Chi-Tin Shih, Yen-Jen Lin, Cheng-Te Wang, Ting-Yuan Wang, Chih-Chen Chen, Ta-Shun Su, Chung-Chuang Lo, and Ann-Shyn Chiang. Diverse community structures in the neuronal-level connectome of the drosophila brain. Neuroinformatics, 18:267–281, 2020.

[43] Maxwell H Turner, Kevin Mann, and Thomas R Clandinin. The connectome predicts resting-state functional connectivity across the drosophila brain. Current Biology, 31(11): 2386–2394, 2021.

[44] Karan KH Manjunatha, Matteo Bruzzone, Giorgio Nicoletti, Samir Suweis, and Marco Dal Maschio. A network-based method for extracting the organization of brain-wide circuits from reconstructed connectome datasets. bioRxiv, pages 2023–05, 2023.

[45] Rodrigo Perin, Thomas K Berger, and Henry Markram. A synaptic organizing principle for cortical neuronal groups. Proceedings of the National Academy of Sciences, 108(13): 5419–5424, 2011.

[46] Nicolas Brunel. Is cortical connectivity optimized for storing information? Nature neuroscience, 19(5):749–755, 2016.

[47] Daniel Witvliet, Ben Mulcahy, James K Mitchell, Yaron Meirovitch, Daniel R Berger, Yuelong Wu, Yufang Liu, Wan Xian Koh, Rajeev Parvathala, Douglas Holmyard, et al. Connectomes across development reveal principles of brain maturation. Nature, 596(7871): 257–261, 2021.

[48] Michael Winding, Benjamin D Pedigo, Christopher L Barnes, Heather G Patsolic, Youngser Park, Tom Kazimiers, Akira Fushiki, Ingrid V Andrade, Avinash Khandelwal, Javier Valdes-Aleman, et al. The connectome of an insect brain. Science, 379(6636):eadd9330, 2023.

[49] MICrONS Consortium, J Alexander Bae, Mahaly Baptiste, Caitlyn A Bishop, Agnes L Bodor, Derrick Brittain, JoAnn Buchanan, Daniel J Bumbarger, Manuel A Castro, Brendan Celii, et al. Functional connectomics spanning multiple areas of mouse visual cortex. bioRxiv, pages 2021–07, 2021. doi: 10.1101/2021.07.28.454025.

[50] Henry Markram, Eilif Muller, Srikanth Ramaswamy, Michael W Reimann, et al. Reconstruction and simulation of neocortical microcircuitry. Cell, 163(2):456–492, 2015.

[51] György Buzsáki and Kenji Mizuseki. The log-dynamic brain: how skewed distributions affect network operations. Nature Reviews Neuroscience, 15(4):264–278, 2014.

[52] Patric Hagmann, Maciej Kurant, Xavier Gigandet, Patrick Thiran, Van J Wedeen, Reto Meuli, and Jean-Philippe Thiran. Mapping human whole-brain structural networks with diffusion mri. PloS one, 2(7):e597, 2007.

[53] Travis A Jarrell, Yi Wang, Adam E Bloniarz, Christopher A Brittin, Meng Xu, J Nichol Thomson, Donna G Albertson, David H Hall, and Scott W Emmons. The connectome of a decision-making neural network. science, 337(6093):437–444, 2012.

[54] Lav R Varshney, Beth L Chen, Eric Paniagua, David H Hall, and Dmitri B Chklovskii. Structural properties of the caenorhabditis elegans neuronal network. PLoS computational biology, 7(2):e1001066, 2011.

[55] Matthew Kahle. Topology of random clique complexes. Discrete mathematics, 309(6): 1658–1671, 2009.

[56] Carina Curto, Chad Giusti, Keler Marku, Eva Pastalkova, and Vladimir Itskov. Pairwise correlation graphs from hippocampal population activity have highly non-random, low-dimensional clique topology. BMC neuroscience, 14(1):1–2, 2013.

[57] Chad Giusti, Eva Pastalkova, Carina Curto, and Vladimir Itskov. Clique topology reveals intrinsic geometric structure in neural correlations. Proceedings of the National Academy of Sciences, 112(44):13455–13460, 2015.

[58] David Eppstein Joseph Wang. Fast approximation of centrality. Graph algorithms and applications, 5(5):39, 2006.

[59] András Ecker, Daniela Egas Santander, Marwan Abdellah, Jorge Blanco Alonso, Sirio Bolaños-Puchet, Giuseppe Chindemi, Dhuruva Priyan Gowri Mariyappan, James B. Isbister, James Gonzalo King, Pramod Kumbhar, Ioannis Magkanaris, Eilif B. Muller, and Michael W. Reimann. Assemblies, synapse clustering and network topology interact with plasticity to explain structure-function relationships of the cortical connectome. bioRxiv, 2023. doi: 10.1101/2023.08.07.552264.

[60] Eyal Gal et al. Rich cell-type-specific network topology in neocortical microcircuitry. Nat. Neurosci., 20(7):1004–1013, 2017.

[61] Michael Okun, Nicholas A Steinmetz, Lee Cossell, M Florencia Iacaruso, Ho Ko, Péter Barthó, Tirin Moore, Sonja B Hofer, Thomas D Mrsic-Flogel, Matteo Carandini, et al. Diverse coupling of neurons to populations in sensory cortex. Nature, 521(7553):511–515, 2015.

[62] Amiram Grinvald, Amos Arieli, Misha Tsodyks, and Tal Kenet. Neuronal assemblies: single cortical neurons are obedient members of a huge orchestra. Biopolymers: Original Research on Biomolecules, 68(3):422–436, 2003.

[63] Michael W Reimann, Henri Riihimäki, Jason P Smith, Jānis Lazovskis, Christoph Pokorny, and Ran Levi. Topology of synaptic connectivity constrains neuronal stimulus representation, predicting two complementary coding strategies. PloS one, 17(1):e0261702, 2022.

[64] Pedro Conceição, Dejan Govc, Jānis Lazovskis, Ran Levi, Henri Riihimäki, and Jason P Smith. An application of neighbourhoods in digraphs to the classification of binary dynamics. Network Neuroscience, 6(2):528–551, 2022.

[65] Hsi-Ping Wang, Donald Spencer, Jean-Marc Fellous, and Terrence J Sejnowski. Synchrony of thalamocortical inputs maximizes cortical reliability. Science, 328(5974):106–109, 2010.

[66] Susanne Schreiber, Jean-Marc Fellous, D Whitmer, Paul Tiesinga, and Terrence J Sejnowski. A new correlation-based measure of spike timing reliability. Neurocomputing, 52: 925–931, 2003.

[67] Christoph Pokorny, Omar Awile, James B Isbister, Kerem Kurban, Matthias Wolf, and Michael W Reimann. A connectome manipulation framework for the systematic and reproducible study of structure–function relationships through simulations. bioRxiv, 2024. doi: 10.1101/2024.05.24.593860.

[68] Kael Dai, Juan Hernando, Yazan N Billeh, Sergey L Gratiy, Judit Planas, Andrew P Davison, Salvador Dura-Bernal, Padraig Gleeson, Adrien Devresse, Benjamin K Dichter, et al. The sonata data format for efficient description of large-scale network models. PLoS computational biology, 16(2):e1007696, 2020.

[69] Andrew T Reid, Drew B Headley, Ravi D Mill, Ruben Sanchez-Romero, Lucina Q Uddin, Daniele Marinazzo, Daniel J Lurie, Pedro A Valdés-Sosa, Stephen José Hanson, Bharat B Biswal, et al. Advancing functional connectivity research from association to causation. Nature neuroscience, 22(11):1751–1760, 2019.

[70] Dmitri B Chklovskii, Thomas Schikorski, and Charles F Stevens. Wiring optimization in cortical circuits. Neuron, 34(3):341–347, 2002.

[71] Georgios Iatropoulos, Wulfram Gerstner, and Johanni Brea. Two-factor synaptic consolidation reconciles robust memory with pruning and homeostatic scaling. bioRxiv, pages 2024–07, 2024.

[72] Claudius Gros. A devil’s advocate view on ‘self-organized’brain criticality. Journal of Physics: Complexity, 2(3):031001, 2021.

[73] Bengier Ülgen Kilic and Dane Taylor. Simplicial cascades are orchestrated by the multidimensional geometry of neuronal complexes. Communications Physics, 5(1):278, 2022.

[74] Max Nolte, Eyal Gal, Henry Markram, and Michael W Reimann. Impact of higher order network structure on emergent cortical activity. Network Neuroscience, 4(1):292–314, 2020.

[75] Rodney J Douglas and Kevan AC Martin. Neuronal circuits of the neocortex. Annu. Rev. Neurosci., 27:419–451, 2004.

[76] Kenneth D Miller. Canonical computations of cerebral cortex. Current opinion in neurobiology, 37:75–84, 2016.

[77] Wolfgang Maass, Thomas Natschläger, and Henry Markram. Real-time computing without stable states: A new framework for neural computation based on perturbations. Neural computation, 14(11):2531–2560, 2002.

[78] Giovanni Petri, Paul Expert, Federico Turkheimer, Robin Carhart-Harris, David Nutt, Peter J Hellyer, and Francesco Vaccarino. Homological scaffolds of brain functional networks. Journal of The Royal Society Interface, 11(101):20140873, 2014.

[79] Vincent Zimmern. Why brain criticality is clinically relevant: a scoping review. Frontiers in neural circuits, 14:565335, 2020.

[80] Inês Hipólito, Jonas Mago, Fernando E Rosas, and Robin Carhart-Harris. Pattern breaking: a complex systems approach to psychedelic medicine. Neuroscience of Consciousness, 2023 (1):niad017, 2023.

[81] Henri Riihimäki. Simplicial q -connectivity of directed graphs with applications to network analysis. SIAM Journal on Mathematics of Data Science, 5(3):800–828, 2023.

[82] Gergő Orbán, Pietro Berkes, József Fiser, and Máté Lengyel. Neural variability and sampling-based probabilistic representations in the visual cortex. Neuron, 92(2):530–543, 2016.

[83] Ruben S Van Bergen and Janneke FM Jehee. Probabilistic representation in human visual cortex reflects uncertainty in serial decisions. Journal of Neuroscience, 39(41):8164–8176, 2019.

[84] Bruno B Averbeck, Peter E Latham, and Alexandre Pouget. Neural correlations, population coding and computation. Nature reviews neuroscience, 7(5):358–366, 2006.

[85] Katherine Morrison and Carina Curto. Predicting neural network dynamics via graphical analysis. In Algebraic and Combinatorial Computational Biology, pages 241–277. Elsevier, 2019.

[86] Caitlyn Parmelee, Samantha Moore, Katherine Morrison, and Carina Curto. Core motifs predict dynamic attractors in combinatorial threshold-linear networks. PloS one, 17(3): e0264456, 2022.

[87] Carina Curto, Christopher Langdon, and Katherine Morrison. Robust motifs of threshold-linear networks. arXiv preprint arXiv:1902.10270, 2019.

[88] Michael W Reimann, Anna-Lena Horlemann, Srikanth Ramaswamy, Eilif B Muller, and Henry Markram. Morphological diversity strongly constrains synaptic connectivity and plasticity. Cerebral Cortex, 27(9):4570–4585, 2017.

[89] Lida Kanari, Ying Shi, Alexis Arnaudon, Natalí Barros-Zulaica, Ruth Benavides-Piccione, Jay S Coggan, Javier DeFelipe, Kathryn Hess, Huib D Mansvelder, Eline J Mertens, et al. Of mice and men: topologically complex dendrites assemble uniquely human networks. bioRxiv, pages 2023–09, 2023.

[90] Michael W Reimann, Sirio Bolanõs-Puchet, Jean-Denis Courcol, Daniela Egas Santander, Alexis Arnaudon, Benoît Coste, Fabien Delalondre, Thomas Delemontex, Adrien Devresse, Hugo Dictus, et al. Modeling and simulation of neocortical micro-and mesocircuitry. Part I: Anatomy. eLife, 13, 2024.

[91] Michael W Reimann, Daniela Egas Santander, András Ecker, and Eilif B Muller. Specific inhibition and disinhibition in the higher-order structure of a cortical connectome. bioRxiv, pages 2023–12, 2023.

[92] Sen Song, Per Jesper Sjöström, Markus Reigl, Sacha Nelson, and Dmitri B Chklovskii. Highly nonrandom features of synaptic connectivity in local cortical circuits. PLoS biology, 3(3):e68, 2005.

[93] Daniel Lütgehetmann, Dejan Govc, Jason P Smith, and Ran Levi. Computing persistent homology of directed flag complexes. Algorithms, 13(1):19, 2020.

[94] Donald B Johnson. Finding all the elementary circuits of a directed graph. SIAM Journal on Computing, 4(1):77–84, 1975.

[95] Louis K Scheffer, C Shan Xu, Michal Januszewski, Zhiyuan Lu, Shin-ya Takemura, Kenneth J Hayworth, Gary B Huang, Kazunori Shinomiya, Jeremy Maitlin-Shepard, Stuart Berg, et al. A connectome and analysis of the adult drosophila central brain. elife, 9: e57443, 2020.

[96] Daniela Egas Santander, Christoph Pokorny, András Ecker, Jānis Lazovskis, Matteo Santoro, Jason P. Smith, Kathryn Hess, Ran Levi, and Michael W. Reimann. Dataset related to “Efficiency and reliability in biological neural network architectures“, April 2024. URL 10.5281/zenodo.10812496.

[97] Michael Reimann, András Ecker, A Sato, Kerem Kurban, and Olli Lupton. Blue-Brain/ConnectomeUtilities: New features and examples: Reciprocal views. 10.5281/zenodo.10059227, October 2023.

[98] M. W. Reimann. Connectomics of (part of) the microns mm3 dataset. 10.5281/zenodo.8364070, September 2023.

[99] Michael Reimann, Daniela Egas Santander, and Christoph Pokorny. Connectivity matrix of the internal connectivity of an earlier version of an SSCX model, January 2024. URL 10.5281/zenodo.10599103.

[100] Pramod Kumbhar, Michael Hines, Jeremy Fouriaux, Aleksandr Ovcharenko, James King, Fabien Delalondre, and Felix Schürmann. CoreNEURON : An Optimized Compute Engine for the NEURON Simulator. Frontiers in Neuroinformatics, 13(63), 2019. doi: 10.3389/fninf.2019.00063.

[101] M. E. J. Newman. The Structure and Function of Complex Networks. SIAM Review, 45 (2):167–256, January 2003. ISSN 0036-1445, 1095-7200. doi: 10.1137/S003614450342480. URL http://epubs.siam.org/doi/10.1137/S003614450342480.

[102] Leonid V Kantorovich. Mathematical methods of organizing and planning production. Management science, 6(4):366–422, 1960.

[103] Catherine S Cutts and Stephen J Eglen. Detecting pairwise correlations in spike trains: an objective comparison of methods and application to the study of retinal waves. Journal of Neuroscience, 34(43):14288–14303, 2014.

[104] Eyal Gal, Rodrigo Perin, Henry Markram, Michael London, and Idan Segev. Neuron geometry underlies universal network features in cortical microcircuits. bioRxiv, 2020. doi: 10.1101/656058.

[105] Daniel Zwillinger and Stephen Kokoska. CRC standard probability and statistics tables and formulae. Crc Press, 1999.

